# Defining the Immunomodulatory Determinants of 3-mer Oligonucleotides on TLR7 and TLR8 sensing

**DOI:** 10.64898/2026.04.29.720527

**Authors:** Sunil Sapkota, Zhikuan Zhang, Christine Wolf, Akiko Fujimura, Daniel S. Wenholz, Mary Speir, Lorna Wilkinson-White, ChakYin Chan, Pauline Percier, Abdallah Ladaycia, W. Samantha N. Jayasekara, Arwaf S. Alharbi, Alexandra L. McAllan, Erandi Rupasinghe, Nathan Beals, Rui Gao, Ruitao Jin, Prashanthi M. Sureshbabu, Marco Westkemper, Liza Cubeddu, Ben Corry, Craig Ramirez, Andrew Hauser, Kim A. Lennox, Mark A. Behlke, Umeharu Ohto, Remi Rosière, Roland Gamsjaeger, Min Ae Lee-Kirsch, Julia I. Ellyard, Olivier F. Laczka, Toshiyuki Shimizu, Michael P. Gantier

## Abstract

Chemical modifications such as 2′-O-methyl (2′-OMe) are central to the efficacy and tolerability of RNA therapeutics. We recently identified that 2′-OMe RNA fragments as short as three nucleotides can exert opposing effects on Toll-like receptor 8 (TLR8) sensing in a motif-dependent manner. This discovery raises important considerations for degradation products of chemically modified RNA therapeutics, which may generate such immunologically active fragments. Here, leveraging their short length, we systematically map how base and sugar modifications within 3-mer oligonucleotides regulate TLR7 and TLR8 responses, and we resolve the structural basis for both TLR8 potentiation and TLR7/8 antagonism by RNA fragments. Building on these insights, we report the development of a dual TLR7/8 inhibitory oligonucleotide with therapeutic potential in autoimmune disease. Together, these findings provide unprecedented resolution of the immunomodulatory properties of oligonucleotide modifications on TLR7/8 and establish 3-mer oligonucleotides as the shortest functional class of RNA therapeutics described to date.

---

Chemical modification is fundamental to oligonucleotide therapeutics, enhancing target engagement, cellular uptake, and nuclease resistance^1, 2^, while also mitigating toxicities such as unintended Toll-like Receptor (TLR)-mediated immune activation^3, 4^. Increasing evidence shows that endosomal TLR7/8/9 primarily sense degradation fragments rather than full length oligonucleotide (oligos), with distinct fragments cooperatively binding different receptor pockets to drive inflammation^5–10^. In this context, our prior analysis of a panel of over 200 2′-O-methyl (2′-OMe) gapmer antisense oligos (ASOs) demonstrated that 75% potentiated TLR8 signalling^11^. Furthermore, short 2′-OMe 3-mer fragments derived from these ASOs could also elicit TLR8 potentiation^12^. Notably, we showed that specific 3-mer motifs with 2′-OMe/DNA modifications can also either potentiate TLR8 or, conversely, block its activation through interaction with an antagonist binding site shared with TLR7, revealing sequence and chemistry dependent determinants of innate immune sensing^12^. This is particularly pertinent in the case of gapmer ASOs, where the central DNA region is flanked by 5′-end modified bases.

Here we show that, beyond 2′-OMe oligos, specific 3-mer oligos bearing 5′-end modifications such as O-methoxyethyl (MOE) and Locked Nucleic Acid (LNA) and a central _d_C moiety can also potentiate TLR8 by engaging its oligomer binding site 2^8^. Building on this observation, we systematically mapped how sugar and base-modifications shape 3-mer-mediated TLR7/8 inhibition and describe the pre-clinical development of a dual TLR7/8 inhibitory 3-mer for the treatment of autoimmunity. Together, these findings expand our understanding of the immunomodulatory activities of oligo degradation fragments on TLR7 and TLR8 and provide design principles for the development of chemically-modified oligos that either harness or circumvent TLR7/8 sensing.

## Results

### TLR8 potentiation by 3-mer oligos is dependent on _m_X_d_C moiety

TLR8 sensing is the result of cooperative binding of RNA degradation products to two sites: one that binds uridine or small-molecule agonists (site 1) and another that engages short uridine-rich fragments, such as _r_U_r_G (where _r_ is RNA) (site 2)^8^. We identified that certain phosphorothioate (PS)-modified 3-mers containing a 5′-end 2′-OMe guanosine and one or two DNA bases, such as _m_G_m_U_d_C or _m_G_d_C_d_C (where _d_ is DNA), can potentiate TLR8 activation by a site 1 agonist, suggesting that these oligos may engage the receptor’s site 2 pocket despite lacking uridine^12^. To extend these observations beyond the case of 5′-end 2′-OMe guanosine, we tested a panel of 16 _m_C_d_X_d_X and 16 _m_U_d_X_d_X oligos on TLR8 sensing. In accord with our analyses of _m_G_d_X_d_X oligos^12^, these experiments revealed a clear preference for potentiation activity for central _d_C moieties, with all eight _m_U_d_C_d_X and _m_C_d_C_d_X oligos being in the top nine best TLR8 potentiators; these hits were independently validated in *TLR7*-deficient THP-1 cells (Fig. 1a, Extended Data Fig. 1a and Supplementary Table S1). A panel of 16 _m_X_d_X and 16 _d_X_d_X 2-mers further highlighted the role for the _d_C moiety in TLR8 potentiation, with the four _m_X_d_C oligos potentiating sensing at 5 μM, with _m_G_d_C being the strongest potentiator (Fig. 1b, Extended Data Fig. 1a and Supplementary Table S1). However, the lack of activity of DNA 3-mers and 2-mers (Fig. 1b, Supplementary Table S1 and ^12^) highlights the crucial contribution of 2′-OMe-modified nucleosides to TLR8 potentiation by _d_C in the context of these short oligos.

**Figure 1.**
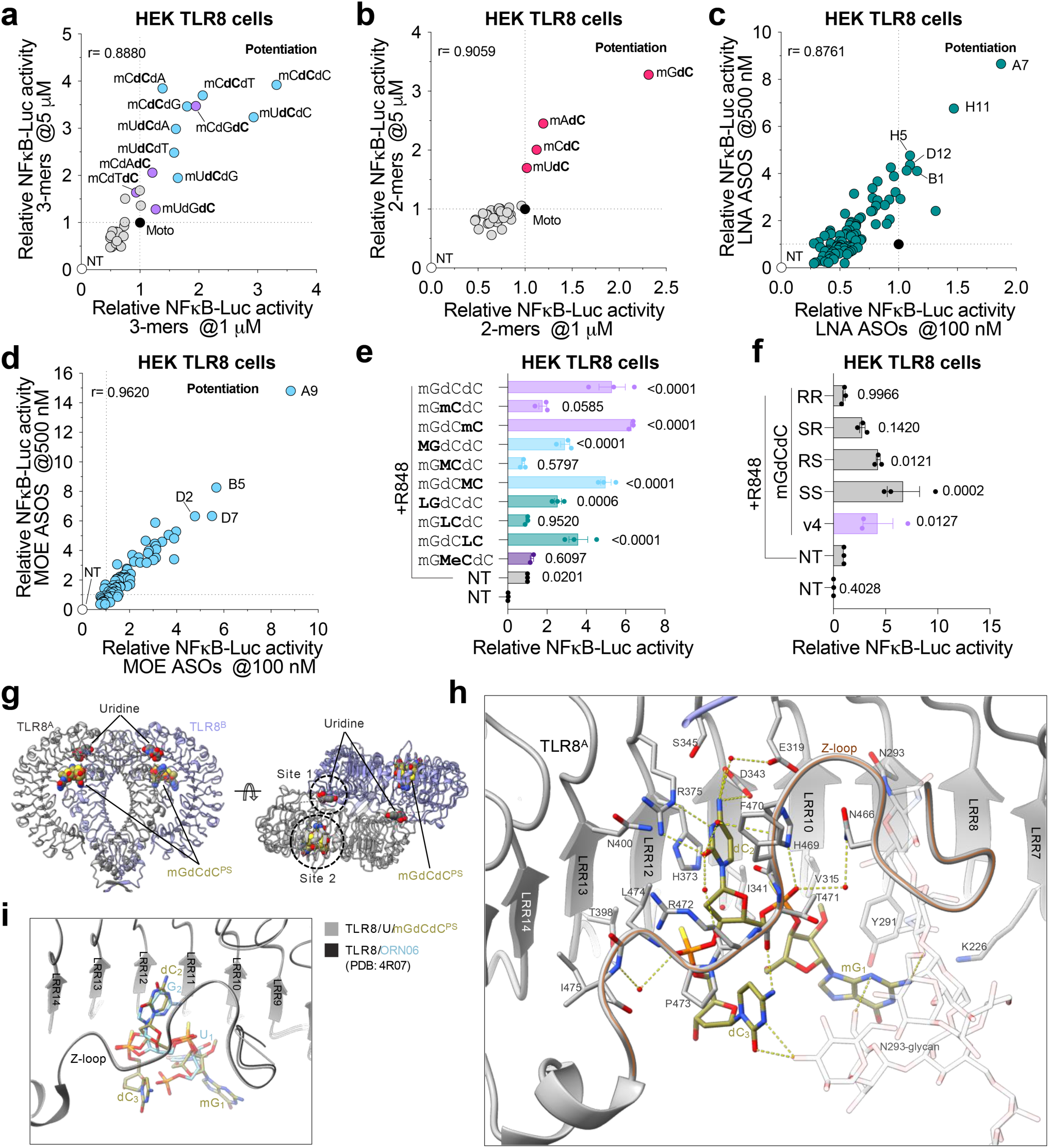
Definition of structural determinants of TLR8 potentiation by 3-mer oligos. (**a-f**) HEK TLR8 cells were pre-treated ∼30 min with 1 μM and 5 μM (**a** and **b**), 100 nM and 500 nM (**c** and **d**), or 5 μM (**e**, **f**) of the indicated oligos prior to overnight stimulation with 600 nM Motolimod (**a** and **b**) and 1 µg/ml R848 (**c** and **d**) followed by luciferase assay. (**a-d**) Data are averaged from 2 or 3 biological replicate for each screen, and the screens at the different oligo concentrations were conducted on independent days. (**e** and **f**) Data are mean of n=3 independent experiments. Data were background-corrected using the non-treated (NT) condition and are shown as relative expression to R848 only (± s.e.m. and one-way ANOVA with uncorrected Fisher’s LSD tests shown compared to NT only (**e**) condition and R848 only (**f**) condition; e and f:*p<0.0001*). (**g**) Crystal structure of the TLR8/U/_m_G_d_C_d_C^PS^ complex. Two TLR8 protomers are shown in cartoon representation. U and _m_G_d_C_d_C^PS^ are shown in sphere representation. (**h**) Close-up view of the _m_G_d_C_d_C^PS^ recognition at the site 2 of TLR8. Residues within 4.5 Å from _m_G_d_C_d_C^PS^ are shown in stick representations. N293 glycan is shown in stick representations. Water molecules are shown using red dots. Sticks are colored by atoms, with the N, O, P and S atoms colored by blue, red, orange and yellow, respectively. Yellow dashed lines indicate hydrogen bonds. (**i**) Structural alignment of TLR8/U/_m_G_d_C_d_C^PS^ and TLR8/ORN06 (PDB: 4R07) complexes at site 2. Structural alignment was performed using the Matchmaker tool in ChimeraX.

To gain a broader insight into the effect of other common gapmer modifications on TLR8 sensing, we screened a panel of 91 LNA and 76 2′-MOE gapmer ASOs on TLR8 sensing, and the top hits were validated in *TLR7*-deficient THP-1 cells (Fig. 1c, 1d, Extended Data Fig. 1b and Supplementary Table S1). Although TLR8 potentiation was not as frequent for these oligonucleotides as previously observed for 2′-OMe ASOs^11^, 50% of the MOE and 27% of the LNA ASOs potentiated TLR8 more than two-fold at 500 nM (Fig. 1c, 1d and Supplementary Table S1), indicating that these chemical modifications of gapmer ASOs still allowed generation of site 2 ligands for TLR8. As with 2′-OMe ASOs, TLR7 inhibition was broadly observed with 85% LNA and 95% MOE ASOs resulting in a 50% reduction in TLR7 sensing at a concentration of 500 nM (Extended Data Fig. 1c, 1d and Supplementary Table S1). Screening 64 MOE 3-mers identified only a single sequence, _M_G_M_G_M_T (where _M_ is MOE), that potentiated TLR8 sensing, in contrast to the 9/64 2′-OMe 3-mers previously observed to potentiate >1.5 fold under the same conditions^12^ (Supplementary Table S1). This suggests that complete modification of 3-mers with MOE impedes binding to TLR8 site 2.

Nonetheless, select substitution of the 5′-end of _m_G_d_C_d_C with MOE and LNA moieties demonstrated that 5′-end MOE and LNA modification were also tolerated for TLR8 potentiation (Fig. 1e). Conversely, sugar modification of the central _d_C of _m_G_d_C_d_C with 2′-OMe, MOE, or LNA, as well as base modification with 5-methylcytidine strongly decreased TLR8 potentiation, confirming the essential role of the _d_C moiety. Stereopure samples of the four PS stereoisomers of _m_G_d_C_d_C (referred to as *RR*, *RS*, *SR*, and *SS*) were also tested. While the *SR*, *RS* and *SS* _m_G_d_C_d_C configurations all retained significant potentiating activity – the *SS* configuration being the strongest – potentiation was lost with _m_G_d_C_d_C-*RR*, suggesting that some structural conformations driven by the PS-backbone are preferred for binding to site 2 (Fig. 1f).

### TLR8 potentiation by 3-mer oligos is dependent on binding to site 2

Our previous description of the broad functional TLR8 potentiation by _m_G_d_C_d_C next led us to interrogate its molecular binding to recombinant human TLR8. We determined the crystal structure of TLR8 in complex with _m_G_d_C_d_C in the presence of uridine (Fig. 1g and Supplementary Table S2). The TLR8/_r_U/_m_G_d_C_d_C complex forms a typical activated dimer structure, in which uridine and _m_G_d_C_d_C bind to site 1 and site 2 of TLR8, respectively (Fig. 1g). Because the _m_G_d_C_d_C used for structure determination contained 4 stereoisomers, we modeled only the most active *SS* isomer for clarity. One _m_G_d_C_d_C molecule is recognized at site 2, which comprises the concave surface of LRR7-13 and the Z-loop located inside a single TLR8 molecule. The middle _d_C_2_ deeply inserts into the site 2 pocket, while _m_G_1_ and _d_C_3_ point outward from site 2 and face the solvent (Fig. 1h). Accordingly, _d_C_2_ is tightly recognized, whereas the recognition of _m_G_1_ and _d_C_3_ is looser. The cytosine base of _d_C_2_ is sandwiched between H373 and H469 side chains. This cytosine base forms multiple direct hydrogen bonds with D343 and R375 side chains, as well as hydrogen bond networks with E319, F470 and R472 via water molecules. The phosphorothioate groups form several hydrogen bonds with the H469 side chain and nearby water molecules. Compared to the previously reported recognition of _r_U_r_G (a degradation product of ORN06 ssRNA) by TLR8^8^, the recognition of _d_C_2_ and the two phosphorothioate groups in _m_G_d_C_d_C resembles that of _r_G_2_ and the phosphate groups in _r_U_r_G, as they superpose well (Fig. 1i). In contrast, the orientations of _m_G_1_ in _m_G_d_C_d_C and _r_U_1_ in _r_U_r_G are opposite. The 2′-OMe group of _m_G_1_ forms characteristic hydrophobic interaction with V315 and I341 side chains. The guanine base of _m_G_1_ contacts the Y291 side chain and a bulky N-glycan attached to N293. The cytosine base of _d_C_3_ forms hydrogen bonds with the T471 main chain O atom and the nearby N293 glycan, an interaction which is not observed in _r_U_r_G recognition. However, the electron densities of _m_G_1_ and the N293 glycan are less well-defined, indicating higher flexibility and loose recognition (Extended Data Fig. 1e). In summary, TLR8 potentiation by _m_G_d_C_d_C is achieved by binding to TLR8 site 2, featuring the tight recognition of the middle _d_C_2_ and the looser recognition of the adjacent moieties.

### Structure activity relationship analyses of TLR7/8 inhibition by 3-mer oligos

Building on our earlier finding that 3′-end DNA substitution in _m_G_m_U_m_C preserves TLR7 inhibition^12^, we systematically evaluated sugar– and base-modified variants at the 3′-end nucleotide by screening at a single concentration (Fig. 2a,b) to further define how sugar and base modifications impact TLR7/8 function. With 2′-OMe-cytidine (_m_G_m_U_m_C) as a starting point, 2′-deoxy, 2′-MOE, and LNA sugar modifications (GUC-v1/v12/v13, respectively) were observed to be well tolerated. Testing at lower concentration showed improved TLR7 inhibiting activity for GUC-v13 but only slight improvement for GUC-v12 (Fig. 2c). In contrast 2′-fluor (GUC-v14) and 2′,3′-dideoxyribose (GUC-v24) sugars modestly reduced potency, while 2′-amino, arabinose and morpholino variants (GUC-v46–48, respectively) ablated antagonistic activity. Substitutions to the cytosine ring of _m_G_m_U_m_C and GUC-v1 methylation at the 5-position increased inhibition potency (GUC-v19/20), whilst a bromo substitution at the same position as well as N4-methylation were only tolerated (GUC-v21/56, respectively). However, both 5-hydroxymethyl and 5-propyne substitutions (GUC-v23/26) abrogated TLR7 inhibition, with the former instead resulting in TLR8 potentiation, similar to GUC-v1.

**Figure 2.**
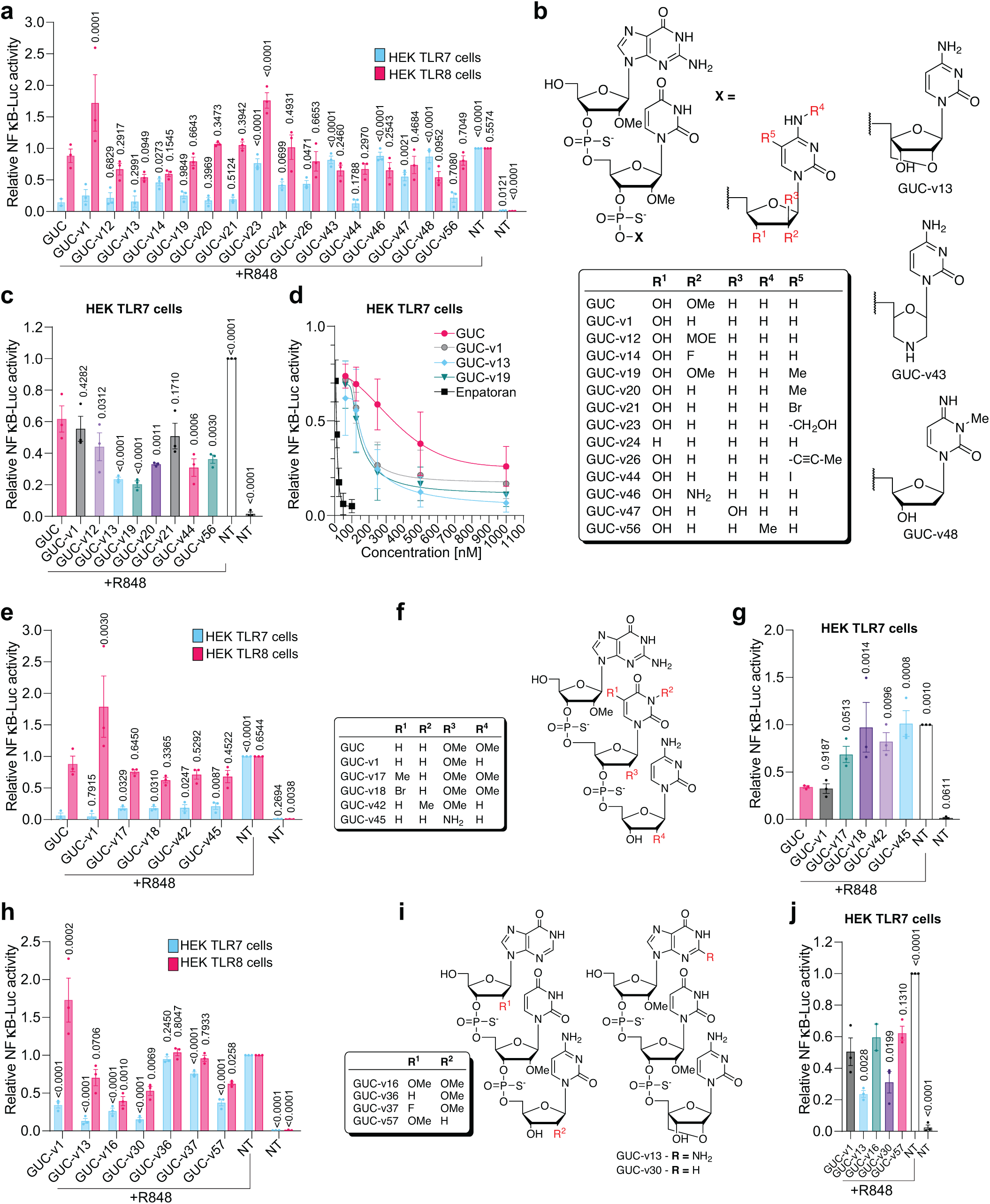
Structure Activity Relationship of TLR7 inhibition and TLR8 potentiation by 3-mer oligos. (**a**, **e** and **h**) HEK-TLR7 and HEK-TLR8 cells were pre-treated ∼30 min with 125 nM (TLR7) and 5 μM (TLR8) of the indicated oligos prior to overnight stimulation with 1 μg/ml of R848 followed by luciferase assay. Data are mean of n=3 independent experiments. Data were background-corrected using the non-treated (NT) condition and are shown as relative expression to R848 only (± s.e.m. and one-way ANOVA with uncorrected Fisher’s LSD tests shown compared to GUC-v1+R848 [**a** (TLR7)], GUC+R848 [**a** (TLR8), and **e**] and R848 [**h**] condition; **a** [TLR7 and TLR8]:*P<0.0001*; **e** [TLR7]: *P<0.0001* and [TLR8]: *P=0.0005*; **h** [TLR7 and TLR8]:*P<0.0001*). (**c**, **g** and **j**) HEK-TLR7 cells were pre-treated ∼30 min with 50 nM of the indicated oligos prior to overnight stimulation with 1 μg/ml of R848 followed by luciferase assay. Data are mean of n=3 independent experiments. Data were background-corrected using the non-treated (NT) condition and are shown as relative expression to R848 only (± s.e.m. and one-way ANOVA with uncorrected Fisher’s LSD tests shown compared to GUC+R848 (**c** and **g**) and GUC-v1+R848 (**j**) condition; **c**, **g** and **j**:*P<0.0001*). (**d**) HEK-TLR7 cells were pre-treated ∼30 min with different doses (1000 nM, 500 nM, 250 nM, 125 nM and 62.5 nM) of the indicated oligos and Enpatoran (100 nM, 50 nM, 25 nM, 12.5 nM and 6.25 nM) prior to overnight stimulation with 1 μg/ml of R848 followed by luciferase assay. Data are mean of n=3 independent experiments. Data were background-corrected using the non-treated (NT) condition and are shown as relative expression to R848 only (± s.e.m. and one-way ANOVA with uncorrected Fisher’s LSD tests shown compared to R848 only condition).

Additional conformationally constrained 2′,4′-bridged chemistries, ENA, cEt, 5-Me-LNA (GUC-v59/60/61, respectively) yielded TLR7 inhibition comparable to GUC-v13 without affecting TLR8 (Extended Data Fig. 2a,b). Dose–response analyses confirmed increased inhibition with GUC-v13 and GUC-v19 over _m_G_m_U_m_C (Fig. 2d), correlating with an ∼3-fold increase in binding affinity strength of GUC-v13 for recombinant *Macaca mulatta* TLR7 (mmTLR7) compared to _m_G_m_U_m_C in isothermal titration calorimetry (ITC) assays (Extended Data Fig. 2c, d). Full replacement of active 3-mer scaffolds with LNA or 2′-MOE abolished TLR7 antagonism (Extended Data Fig. 2e), while screening _m_G_m_X_L_X and _m_U_m_X_L_X libraries still identified _m_G_m_U_L_C (GUC-v13) as the most active inhibitor (Extended Data Fig. 2f). In agreement with the observations discussed earlier on _m_X_d_C, TLR8 potentiation by GUC-v1 was significantly impacted by most modifications to the deoxycytidine, barring 5-hydroxymethyl. We next assessed modifications to the uridine nucleoside (Fig. 2e–g). Modification of the sugar with a 2′-amino group (GUC-v45) and all substitutions to the uracil ring of GUC, including 5-methyl, 5-bromo, N3-methyl (GUC-v17/18/42, respectively), resulted in a mild, but significant, reduction in the potency of TLR7 inhibition (Fig. 2e-g).

Finally, we tested modification of the 5′-end guanosine, the principal TLR7-engaging residue^12^. Conversion of guanosine to inosine in _m_G_m_U_m_C through removal of the guanine base 2-amino group (GUC-v16) preserved TLR7 antagonism, whereas further modifying the 2′-OMe group to 2′-deoxy or 2′-fluor (GUC-v36/37, respectively) was not tolerated (Fig. 2h–j). This suggested that the hydrogen bonding previously reported through the 2′-OMe-G 2-amino group to V355 is non-essential to human TLR7 antagonism^12^. Combining 5′-end 2′-OMe-inosine with a 3′-end 2′-deoxycytidine (GUC-v57) was tolerated whilst LNA modification of the cytidine (GUC-v30) resulted in TLR7 inhibition comparable to GUC-v13. Notably, 5′ 2′-OMe-inosine (GUC-v16/30/57) resulted in an increase in TLR8 inhibition. In summary, 3′-end constrained nucleosides, such as LNA, can enhance TLR7 inhibition, while 5′-end inosine confers dual TLR7/8 inhibitory activity to GUC analogous oligos.

### 5′-end 2′-OMe inosine 3-mers exhibit increased affinity to TLR8 antagonist binding site

To probe the molecular basis of the enhanced TLR8 inhibition conferred by 5′-end 2′-OMe inosine, we investigated the incorporation of this base into the best known TLR8 selective inhibitor _m_G_m_A_d_G^12^. In cell-based assays, _m_I_m_A_d_G displayed markedly improved TLR8 inhibition compared with _m_G_m_A_m_G and _m_G_m_A_d_G (Extended Data Fig. 3a). Surface plasmon resonance (SPR) analyses confirmed that _m_I_m_A_d_G also exhibited a three-fold higher binding affinity for recombinant human TLR8 (Kd ∼875 nM) relative to _m_G_m_A_d_G, whereas GUC-v16 bound more weakly (Kd ∼13 µM), consistent with the negligible affinity of the parent _m_G_m_U_m_C scaffold^12^ and that of GUC-v13 (Fig. 3a and Supplementary Table S3a). ITC analysis of the GUC-v16/mmTLR7 complexation confirmed that the binding of GUC-v16 was ∼1.5-fold stronger than that of parental _m_G_m_U_m_C (Extended Data Fig. 2c,d). Mutational analyses further demonstrated that the binding of GUC-v16 to both recombinant mmTLR7 and hTLR8 was nearly abolished in their respective F507S and F495S antagonist-site mutants^12^, while the binding of resiquimod (TLR7 site 1 ligand) and GCC-v4 (TLR8 site 2 ligand – Fig. 1) remained unaffected (Fig. 3b,3c and Supplementary Table S3b, S3c). This directly confirms that these 3-mer variants engage the conserved antagonist pocket rather than canonical agonist-binding sites.

**Figure 3.**
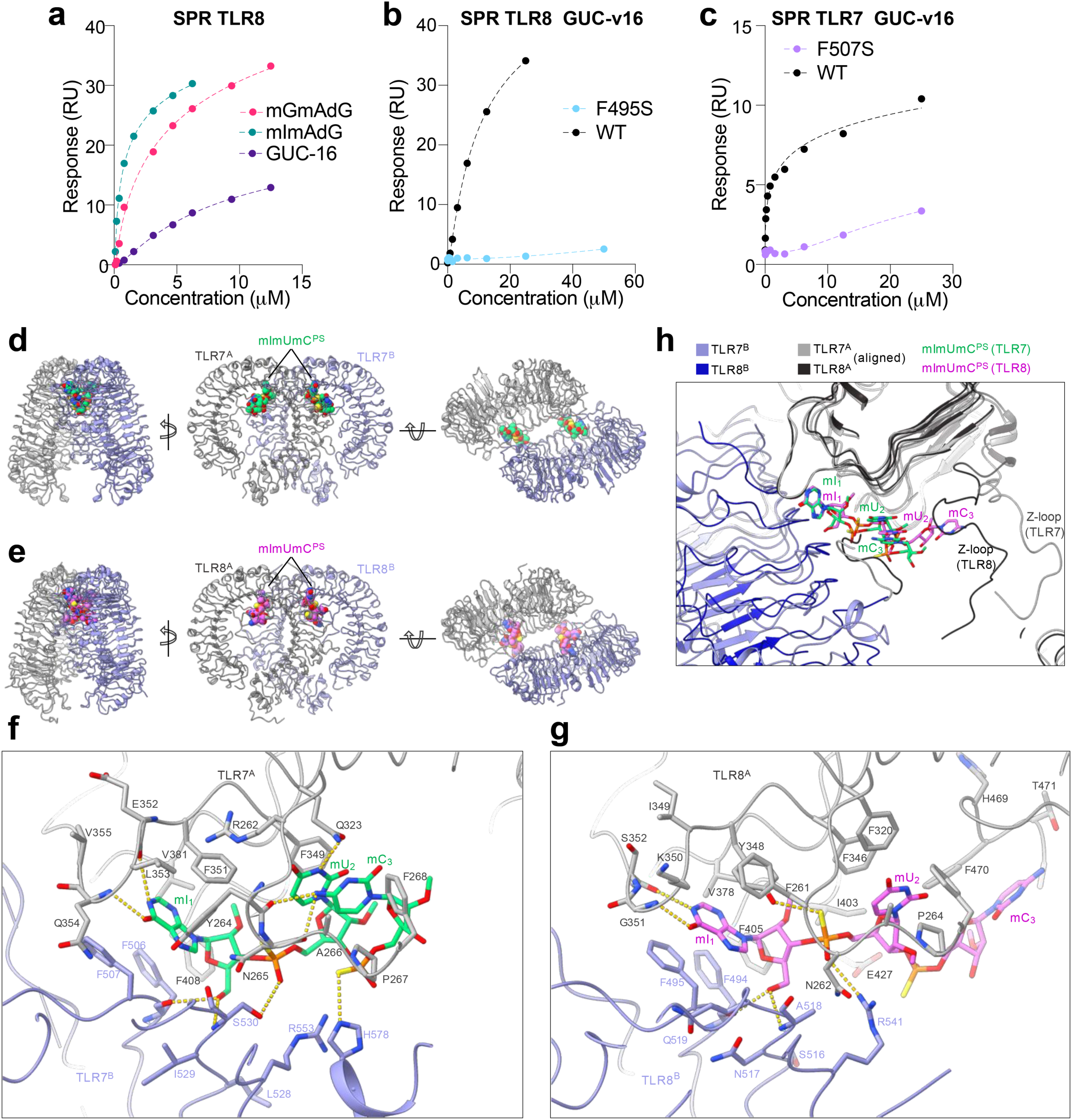
5’-end inosine modification increases 3-mer binding to TLR7/8 antagonist site. (**a-c**) Surface plasmon resonance (SPR) analyses of recombinant wild type human TLR8 (**a**, **b**), wild type mmTLR7 (**c**), mutant TLR8 (F495S) or mutant mmTLR7 (F507S) with the indicated concentrations of indicated 3-mers. Data shown are representative of 3-4 independent analyses (Supplementary Table S3). (**d, e**) Cryo-EM structure of the TLR7/GUC-v16 complex and crystal structure of the TLR8/GUC-v16 complex. Two TLR7 or TLR8 protomers and GUC-v16 are shown in cartoon and sphere representations, respectively. (**f, g**) Close-up views of the GUC-v16 (mImUmC^PS^) recognition at the antagonistic sites of TLR7 and TLR8. Residues within 4.5 Å from GUC-v16 are shown in stick representations. Sticks are colored by atoms, with the N, O, P and S atoms colored by blue, red, orange and yellow, respectively. Yellow dashed lines indicate hydrogen bonds (cutoff distance < 3.5 Å). (**h**) Structural alignment of TLR7/GUC-v16 and TLR8/GUC-v16 complexes. Structural alignment was performed using the Matchmaker tool in ChimeraX.

These findings prompted us to define how GUC-v16 engages the antagonist binding sites of TLR7 and TLR8. We determined the cryo-EM structure of the TLR7–GUC-v16 complex (3.0 Å) and the crystal structure of the TLR8–GUC-v16 complex (3.1 Å), both adopting open, inactive dimers akin to TLR7/8 bound to small molecule antagonists (Methods, Table S2 and Table S4). Because the *RR* stereoisomer was the most potent TLR7 inhibitor in cells (Extended Data Fig. 3b,c), we modelled this isomer in both structures (Fig. 3d-e and Extended Data Fig. 3d,e). Similar to GUC-v1 binding to TLR7^12^, the 5′-end 2′-OMe inosine (mI_1_) sits deep in the antagonist binding site at the protomer interface, its hypoxanthine ring undergoing an aromatic stacking complex with Y264^A^, F351^A^, L353^A^, V381^A^, F408^A^, F506^B^, and F507^B^ and hydrogen bonded via N1–E352 and C6=O–Q354 (Fig. 3f); the 2′-OMe interacts with the hydrophobic pocket formed by F351^A^, V381^A^ and F408^A^ side chains and the 5′-OH group hydrogen-bonds to the F506^B^/S530^B^ backbone. The mU_2_ and mC_3_ moieties are more loosely recognized, solvent-exposed at the ribose without making apparent contact with TLR7, and the phosphorothioates are close to the cationic R553^B^ and H578^B^, forming weak electrostatic interactions (Fig. 3f).

The recognition of GUC-v16 by TLR8 shares similarities with TLR7 but also shows large variations (Fig. 3e-g). The 5′-end mI_1_ is recognized in a highly similar manner: (1) The hypoxanthine moiety is complexed by homologous aromatic and hydrophobic residues in TLR8 including F261^A^, Y348^A^, V378^A^, F405^A^, F494^B^ and F495^B^; (2) the hypoxanthine ring also contacts and forms a hydrogen bond via N1–S352 and C6=O–G351^A^; (3) the interactions between TLR8 and the 2′-OMe and 5′-OH groups of mI_1_ are similar to those observed in TLR7, and TLR8 R541^B^ (corresponding to R553^B^ in TLR7) interacts with the PS groups through electrostatic interactions. Structural alignment of the antagonist binding sites in TLR7 and TLR8 shows that the mI_1_ moieties of GUC-v16 superimpose well, whereas the mU_2_ and mC_3_ adopt a distinct extended conformation that lacks stacking between the pyrimidine bases (Fig. 3h). Interestingly, the mC_3_ partially interacts with a portion of the Z-loop region of TLR8, including residues H469^A^, F470^A^ and S471^A^, which overlaps with site 2 for short RNA binding in the active form of TLR8^8^ (Fig. 3g). However, the electron densities of the mC_3_ moiety and nearby TLR8 residues are weak and less well-defined, indicating higher flexibility in this region (Extended Data Fig.3e). By contrast, the Z-loop region of TLR7 is apart from the antagonistic site and does not contribute to the formation of this site (Fig. 3h).

### GUC-v16 3-mer to limit aberrant TLR7/8 function in disease contexts

Structural analyses relying on a homology model of the GUC-v16 in complex with the mouse (m)TLR7 suggested that the 2′-OMe inosine of GUC-v16 more stably engages the murine TLR7 antagonist binding site than _m_G_m_U_m_C due to additional water-mediated hydrogen-bond network contacts between the inosine base and V381 (Extended Data Fig. 4a). Consistent with this, cell assays showed GUC-v16 inhibits mTLR7 more effectively than _m_G_m_U_m_C, although _m_G_m_G_d_C (GGC-v1) remained the top antagonist (Extended Data Fig. 4b). In *Tlr7^Y264H^* primary bone marrow-derived macrophages (BMDMs) which exhibit basal TLR7 engagement^12^, GUC-v16 and the small molecule TLR7 inhibitor BMS905^13^ suppressed constitutive TLR7 signalling and reduced expression of a defined TLR7-gene set, mirroring the effects observed with Enpatoran and GGC-v1^12^(Fig. 4a; Extended Data Fig. 4c,d). Accordingly, RT-qPCR analyses validated that 7 out of 8 genes significantly down-regulated in RNA-sequencing analyses of BMS905-treated *Tlr7^Y264H^*BMDMs were also significantly down-regulated by GUC-v16 (Fig. 4a; Extended Data Fig. 4c,d). Prophylactic intravenous injection of GUC-v16 complexed with in vivo-jetPEI® blunted R848-induced splenic expression of TLR7-induced genes (*Fpr1*, *Fpr2*, *Marco*, *Nfkbiz* and *TNF*) (Fig. 4b). Similarly, pre-treatment of mice with GUC-v16 formulated as a topical 2.5% cream abolished Aldara-driven skin inflammation without affecting splenomegaly, indicating localized TLR7 inhibition (Fig. 4c; Extended Data Fig. 4e,f). The inhibitory effect of GUC-v16 on R848-driven TLR7 signalling was also robustly observed across many cytokines (e.g. TNF, MIP2, CXCL1) in bronchial alveolar lavages and plasma following the prophylactic intratracheal administration of naked GUC-v16 (Fig. 4d-f). However, GUC-v16 treatment did not significantly impact immune cellular recruitment to the lung (Fig. 4d).

**Figure 4.**
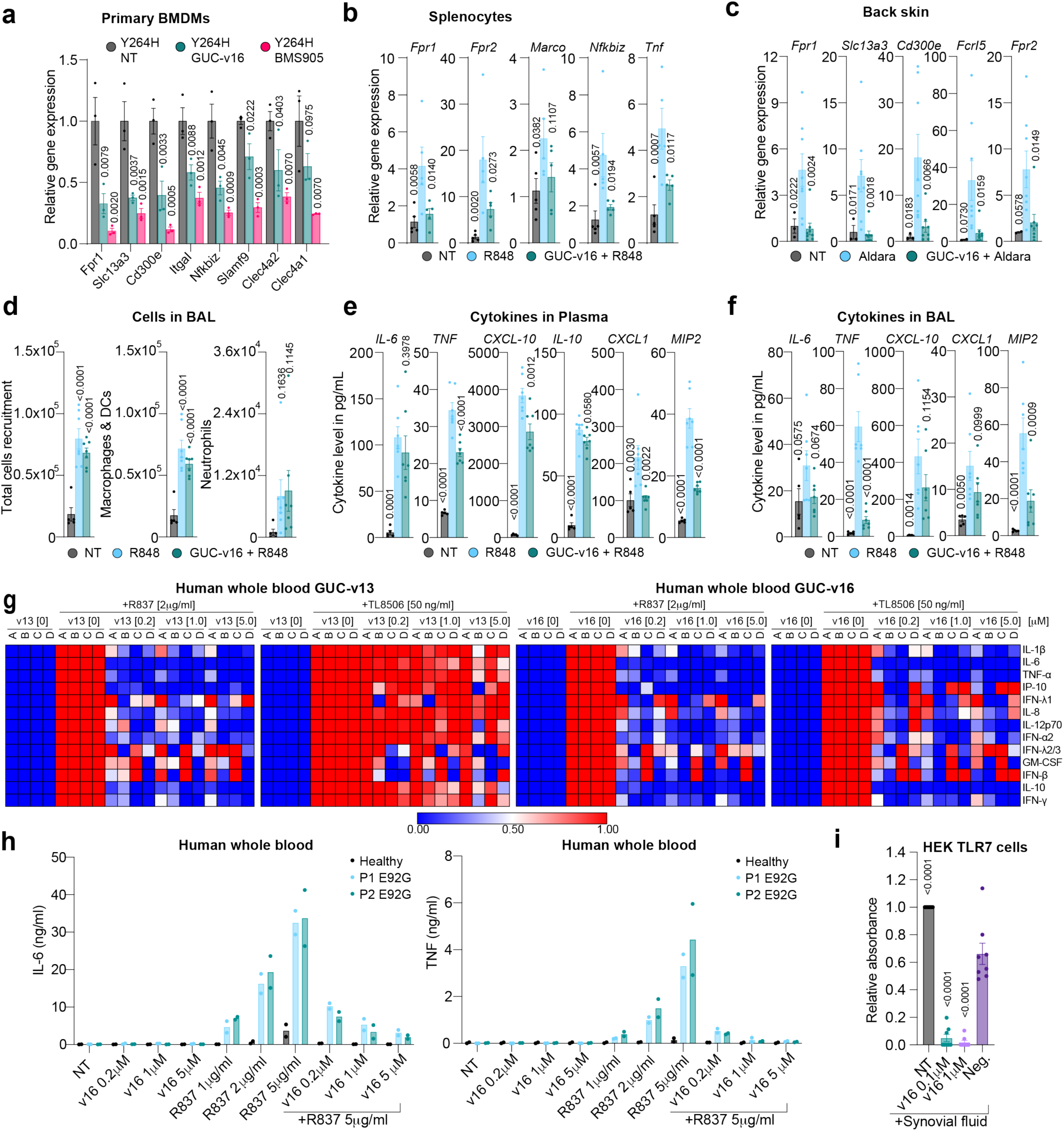
GUC-v16 inhibits aberrant TLR7/8 function in vitro and in vivo. (**a**) Bone-marrow-derived macrophages (BMDMs) from *Tlr7^Y264H^* mice were treated for 24 h with 1 μM of BMS905 and 5 μM of GUC-v16 oligo prior to RNA purification for RNA-sequencing or RT-qPCR analyses. RT-qPCR analyses of *Fpr1, Slc13a3, Cd300e, Itgal, Nfkbiz, Slamf9, Clec4a2* and *Clec4a1* reported to *18S* in RNA lysates from primary BMDMs from 3 independent *Tlr7^Y264H^* mice. Data are shown relative to non-treated (NT) mice (± s.e.m. and two-way ANOVA with uncorrected Fisher’s LSD tests shown compared to NT *Tlr7^Y264H^* condition). (**b**) WT C57/BL6 mice were injected i.v. with 200 μg of GUC-v16 conjugated with *in vivo*-jetPEI® or vehicle (glucose solution) for 1 h prior to i.p. injection of 25 μg R848 for 2 h before collection of spleens. RT-qPCR analyses of *Fpr1, Fpr2, Marco, Nfkbiz* and *Tnf* reported to *Gapdh*, from spleen lysates; data are shown relative to non-treated (NT) mice (± s.e.m. and two-way ANOVA with uncorrected Fisher’s LSD tests shown compared to R848 mice). Each dot represents an individual mouse with bars showing the mean of n=5 mice/group is shown (± s.e.m. and one-way ANOVA with uncorrected Fisher’s LSD tests shown compared to R848 group). (**c**) Aldara cream was applied topically to the back of WT C57/BL6 mice directly following, or not, application of 100 μl of highly pure 2.5% GUC-v16 cream oligonucleotide (>99.4%). After four days, mice were humanely euthanised and back skin collected, and lysed for RNA purification. RT-qPCR analyses of indicated genes reported to that of *18S* expression, relative to NT mice. Data are representative of 2 independent experiments. Mean of n=3 NT and n=8 Aldara/Aldara+GUC-v16 mice/group is shown (± s.e.m. and one-way ANOVA with uncorrected Fisher’s LSD tests shown compared to Aldara group). (**d-f**) Wild type C57BL/6J mice were treated i.t. with 2.5 μg of naked GUC-v16 in water for 1 h prior to. i.t. injection of 50 μg of R848. BALF (**d, f**) were harvested 7 h post R848 injection and analysed by cytospin differential cell counting for total cell counts (**d**) and using an MSD multiplex assay for cytokine quantification (**f**). Cytokine levels in plasma were also quantified using an MSD multiplex assay (**e**). Data shown are from 1 experiment, representative of n=2 independent experiments. Mean of n=5 NT and n=7 R848/R848+GUC-v16 mice/group is shown (± s.e.m. and one-way ANOVA with uncorrected Fisher’s LSD tests shown compared to NT [d] or R848 [**e,f**] group). Whole blood from 4 (**g**) or 1 (**h**) healthy controls or patients with UNC93B1 E92G mutation (**h**) was pre-treated with indicated 3-mer concentration for 30 min prior to stimulation with indicated amount of R837 or 50ng/ml TL8-506 for 24 h prior to cytokine bead analyses. (**g**) For each control, cytokine levels were averaged for each technical replicate (6/sample), background corrected to unstimulated samples only, and reported to cytokine levels from stimulation only controls. Heat maps were generated using scale from 0-1 (see methods), were 1 is the cytokine level of the agonist only control. (**g-h**) Data shown are from a minimum of two independent experiments – conducted on two independent days. (**i**) HEK-TLR7 cells were pre-treated with indicated concentration of oligo for 1 h prior to overnight stimulation with 10% of synovial fluid from RA patient. SEAP absorbances were background-corrected using the non-treated condition, and are shown as relative expression to the synovial fluid only control; Neg. is a non-oligo condition used as control. Data shown are averaged from synovial fluid stimulations from n=8 patients (± s.e.m. and two-way ANOVA with uncorrected Fisher’s LSD tests shown compared to Neg condition).

In human whole blood from four healthy donors, GUC-v16 potently suppressed TLR7/8-driven cytokine production induced by TLR7 and TLR8 agonists (Fig. 4g). In contrast, GUC-v13 inhibited TLR7 (cytokine heatmap with R837) but not TLR8 (cytokine heatmap with TL8-506) (Fig. 4g), and _m_I_m_A_d_G showed dual TLR7/8 inhibiting activity with lower potency than GUC-v16 (Fig. 4g; Extended Data Fig. 4h). In SLE patient blood carrying the *UNC93B1^E92G^* mutation, which displays hyper-responsiveness to R837^14^, GUC-v16 markedly reduced R837-induced IL-6 and TNF (Fig. 4h). Finally, GUC-v16 significantly decreased nuclear factor-kappa B (NF-κB) responses to the synovial fluid from 8 rheumatoid arthritis (RA) patients in HEK-TLR7-expressing cells (Fig. 4i), implying these also contain immunostimulatory RNA products^15^. Collectively, these findings establish that GUC-v16 is a robust dual antagonist of TLR7 and TLR8 in murine models of inflammation, human whole blood, and samples derived from SLE and RA patients.

## Discussion

Predicting toxicities of oligonucleotide therapeutics remains a major challenge for the field. Although adverse responses have historically been attributed to TLR9 sensing of ASOs^3^, emerging evidence indicates that TLR7/8 also contribute to innate immune activation, as recently suggested for the MOE-ASO bepirovirsen^16^. Our work extends this view by showing that full-length MOE– and LNA-modified gapmers can modulate TLR7 (inhibition) and TLR8 (potentiation), though less frequently than 2′-OMe ASOs^11^. Importantly, we demonstrate that 3-mer degradation fragments containing mixed 2′-OMe/2′-MOE or LNA with DNA can potentiate TLR8, and our structural analyses reveal that _d_C-containing motifs with a preceding modified ribose engage TLR8 site 2, supported by a 5′-end 2′-O-Me guanosine, stabilizing the active conformation. These findings confirm that non-RNA oligo fragments can synergize with uridine bound at site 1, consistent with prior reports showing cooperative TLR8 activation by uridine with poly(_d_T) DNA or _d_T-rich ASOs^11, 12, 17, 18^. Given that TLR8 activation reflects competition between agonists and natural antagonist fragments, TLR8-potentiating ASOs may lower tolerance to self and promote inflammation^12^. Although _m_G_d_C_d_C provides a structural exemplar, our broader 3-mer screens reveal that many short motifs can potentiate TLR8, underscoring the need to incorporate TLR8-avoidance principles into ASO or select anti-microRNAs design. Notably, 5-methylcytidine blunted TLR8 potentiation, suggesting that similar to its inhibitory effect on TLR9 sensing of CpG motifs^19^, it could be used to prevent TLR8 potentiation.

In contrast, we show that most LNA– and MOE-gapmer ASOs inhibit TLR7, similar to our observations with 2′-OMe ASOs^11^. This highlights the frequent off-target immune-suppression of TLR7 by this class of oligonucleotides, and the risk this could have for intercurrent infections such as SARS-CoV-2^20^. These data align with our earlier demonstration that poly(_d_T) DNA interacts directly with the antagonistic site of TLR7 in a length-dependent manner, suggesting that sequence-dependent TLR7 inhibition becomes more permissive with longer oligos^12^. Building on our finding that 5′-end 2′-OMe guanosine directs binding to the TLR7 antagonistic site, our SAR analyses of GUC 3-mers show that diverse chemical modifications to cytidine, including 3′-end–constrained nucleosides such as LNA, retain TLR7-antagonistic activity. Nonetheless, fully MOE– or LNA-modified GUX 3-mers lose this activity, suggesting that substituting 2′-OMe guanosine at the 5′ position disrupts antagonism. In the context of gapmers, these results implicate central DNA regions and DNA/MOE or DNA/LNA junctions as the principal sources of TLR7 inhibitory fragments.

We further show that 2′-OMe 3-mers can directly engage the TLR8 antagonist binding site, through critical aromatic stacking interactions between the 5′-end nucleobase and F495. As evidenced with GAG 3-mer analogues, 5′-end mutation with 2′-OMe inosine robustly increased affinity to TLR8, facilitating the development of GUC-v16 as a dual inhibitor of TLR7 and TLR8; 5′-end 2′-OMe inosine substitution of _m_G_m_U_m_C also enhanced activity on mouse TLR7. Accordingly, prophylactic administration of GUC-v16 protects against ligand-induced TLR7 signalling in mice and in whole human blood, and suppresses constitutive TLR7 activation in *Tlr7^Y264H^* mutant cells. Its capacity to block aberrant TLR7/8 sensing in the context of UNC93B1 mutation, and to reduce TLR7-driven inflammation in RA synovial fluid, further supports the potential of such 3-mers as clinical immunomodulators for autoimmune and autoinflammatory diseases.

In summary, we reveal how chemically-modified oligonucleotides and their 3-mer fragments modulate TLR7 and TLR8. While this model depends on nuclease driven fragment generation, itself shaped by chemical modifications, our findings provide a molecular basis for designing safer oligonucleotides that minimize TLR8 activation and TLR7 inhibition.

## Methods

### Cell culture and reagents

293XL-hTLR7 (and HEK-Blue™ TLR7 for Fig. 4i only), 293XL-hTLR8 stably expressing human TLR7 and TLR8, respectively, were purchased from Invivogen and were maintained in Dulbecco’s modified Eagle’s medium plus L-glutamine supplemented with 1× antibiotic/antimycotic (Thermo Fisher Scientific) and 10% heat-inactivated foetal bovine serum (referred to as complete DMEM), with 10-30 μg/ml Blasticidin (Invivogen). WT and TLR7-deficient Human acute myeloid leukemia THP-1 cells^21^ were grown in RPMI 1640 plus L-glutamine medium (Life Technologies) complemented with 1x antibiotic/antimycotic and 10% heat inactivated foetal bovine serum (referred to as complete RPMI). THP-1 cells were not differentiated with PMA unless otherwise noted. RAW264.7-ELAM macrophages^22^ were grown in complete DMEM. All the cells were cultured at 37°C with 5% CO_2_. Cell lines were passaged 2-3 times a week and tested for mycoplasma contamination on routine basis using Mycostrip (Invivogen).

Cells were treated with indicated concentration of oligonucleotides or BMS905 (MedChemExpress) for 20-60 min, prior to R848 (Cayman Chemical), Motolimod (MedChemExpress), R837 (Invivogen), TL8-506 (Invivogen), as indicated. Trimer and longer oligonucleotides were synthesised by Integrated DNA Technologies, Syngenis Pty Ltd or Wuxi AppTec, and resuspended in Rnase-free TE buffer, pH 8.0 (Thermo Fisher Scientific). For *in vivo* experiments, the oligonucleotides were HPLC-purified and confirmed to be endotoxin free by Limulus Amebocyte Lysate gel-clot method. Sequences and modifications are provided in Supplementary Table S1. 2′-MOE is MX, 2′-OMe is mX, DNA is dX (2 and 3-mer) or UPPERCASE alone (16 and 20-mers), RNA is rX, LNA is lX, and phosphorothioate inter-nucleotide linkages are denoted with a *. All the oligos used in this study are fully phosphorothioate modified. A7-LNA-dT: lA*lA*lC* dT*dT*dT* dT*dT*dT* dT*dT*dT* dT*lA*lT* lC.

*Tlr7^Y264H^* C57BL/6NCrl mice (used under Australian National University animal ethics, A2023/309) have a mutation leading to constitutive activation of TLR7 and SLE-like disease^23^. Primary bone marrow derived macrophages (BMDMs) from 6-8 week-old *Tlr7^Y264H^* heterozygous female mice were extracted and differentiated for 5 days in complete DMEM supplemented with L929 conditioned medium as previously reported ^24^, prior to 24 h incubation with 5 μM GUC-v16 or 1 μM BMS905 and total RNA purification for RNA sequencing and RT-qPCRs.

### Study Participants

Whole blood samples from healthy controls included both male and female donors. For analyses of control responses, sex was not evaluated as a biological variable. In comparative analyses between the patient (P1, P2, described in Wolf et al.^14^) and healthy controls, samples were matched for sex and age. The study was approved by the ethics committee of the Medical Faculty, Technische Universität Dresden (BO-EK-446102022), and conducted in accordance with the Declaration of Helsinki.

### Whole blood assays healthy controls and UNC93B1 patients

Heparinized blood was aliquoted (140 µl/well) into 96-well plates, mixed with RPMI medium (Gibco, 31870-025) and maintained at 37°C. For cytokine induction, samples were stimulated with indicated concentrations of R837 (TLR7 agonist; InvivoGen, tlrl-imqs-1) or TLR8-506 (TLR8 agonist, tlrl-tl8506) for 24 hours at 37°C. 3-mer concentration testing involved 30-minute pretreatment at indicated concentrations prior to TLR7 or TLR8 stimulation. Plates were centrifuged (800 ×g, 5 min, RT), and supernatants were transferred to new plates without disturbing cellular layers, then flash-frozen at –80°C.

### Luciferase assays

HEK293 cells stably expressing hTLR8 or hTLR7 were reverse-transfected with pNF-κB-Luc4 reporter (Clontech), with Lipofectamine 2000 (Thermo Fisher Scientific), according to the manufacturer’s protocol. Briefly, 500,000-700,000 cells were reverse-transfected with 200-400 ng of pNF-κB-Luc4 reporter with 1.2 μl of Lipofectamine 2000 per well of a 6-well plate and incubated for 3-24 h at 37 °C with 5% CO_2_. Following transfection, the cells were collected from the 6-wells and aliquoted into 96-wells, just before oligo and overnight TLR stimulation. Similarly, the RAW264.7 cells stably expressing an ELAM-Luc reporter were treated overnight. In all cases, the cells were lysed in 40 μl (for a 96-well plate) of 1X Glo Lysis buffer (Promega) for 10 min at room temperature. 15 μl of the lysate was then subjected to firefly luciferase assay using 35 μl of Luciferase Assay Reagent (Promega). Luminescence was quantified with a Fluostar OPTIMA (BMG LABTECH) luminometer.

## RA Synovial fluid assays

Leftover fully de-identified synovial fluid samples were collected from the knee or hip joints of rheumatoid arthritis patients undergoing routine arthrocentesis at Lahey Hospital & Medical Center. As leftover samples from routine testing these samples were not subject to IRB review and consent procedures. Arthrocentesis, where a ∼18– to 25-gauge is inserted into the joint space and synovial fluid is aspirated into a syringe, was used to collect synovial fluid, and performed under sterile conditions to minimize the risk of infection. HEK-Blue™ TLR7 (Invivogen, hkb-htlr7v2) reporter cells were cultured according to the manufacturer’s instructions and plated in complete DMEM prior to treatment. Cells were pretreated for 1 hour with vehicle or the indicated oligonucleotide, followed by stimulation with synovial fluid obtained from rheumatoid arthritis patients. Synovial fluid was added to achieve a final concentration of 10% (v/v) in each well (15 μl synovial fluid + 135 μl culture medium) and incubated for 24 hours at 37°C. Following stimulation, reporter activity was assessed by adding QUANTI-Blue™ reagent (invivogen, rep-qbs) and incubating at room temperature for 15 minutes. Absorbance was then measured at 620 nm using a GloMax Plate Reader (promega, GM3000). All conditions were performed in triplicate, and blank media controls were included for background subtraction.

### Cytokine analyses

Production of human IP-10 levels were measured in supernatants from THP-1 cells using the IP-10 (BD Biosciences, #550926) ELISA kit. Tetramethylbenzidine substrate (Thermo Fisher Scientific) was used for quantification of the cytokines on a Fluostar OPTIMA (BMG LABTECH) plate-reader. All ELISAs were performed according to the manufacturers’ instructions. Supernatants of human whole blood assays were analyzed using LEGENDplex™ Human Inflammation Panel 1 (BioLegend) per manufacturer protocols. For data acquisition, an LSR Fortessa (BD Biosciences) was used. Data analysis was performed via LEGENDplex™ Data Analysis Suite v8.1 (BioLegend).

### RNA and RT-qPCR analyses

Total RNA was purified from *Tlr7^Y264H^* primary BMDMs or mouse skin biospies using the PureLink RNA Mini Kit (Thermo Fisher Scientific) and DNase-treated using the Purelink DNASE set (Thermo Fisher Scientific). Random hexamer cDNA was synthesised from isolated RNA using the High-Capacity cDNA Archive kits (Thermo Fisher Scientific) according to the manufacturer’s instructions. RT-qPCR was carried out with the Power SYBR Green Master Mix (Thermo Fisher Scientific) on a QuantStudio 6 Flex RT-PCR system (Thermo Fisher Scientific) with the QuantStudio™ Real-Time PCR Software v1.7.2. Each PCR was performed in technical duplicate and mouse and human *18S* (**4a, 4c**) or *Gapdh* (**4b**) were used as the reference genes. Each amplicon was gel-purified and used to generate a standard curve for the quantification of gene expression. Melting curves were used in each run to confirm specificity of amplification (**4a, 4c**). For **4b**, relative gene expression was calculated using the 2^-ΔΔCt^ method. Primers used were the following; Mouse 18s: Rn18s-FWD TAA CCC GTT GAA CCC CAT T; Rn18s-REV CCA TCC AAT CGG TAG TAG CG; M-F-Slc13a3 GGA AGG CCG ATG CCT CTA TG; M-R-Slc13a3 GGA AGT TGG TGT CGA GGA AGT; M-F-Itgal CCA GAC TTT TGC TAC TGG GAC; M-R-Itgal GCT TGT TCG GCA GTG ATA GAG; M-F-Fpr1 CAT TTG GTT GGT TCA TGT GCA A; M-R-Fpr1 AAT ACA GCG GTC CAG TGC AAT; M-F-Fpr2 GAG CCT GGC TAG GAA GGT G; M-R-Fpr2 TGC TGA AAC CAA TAA GGA ACC TG; M-F-Cd300e TGG GTC TTA CTG GTG CAA GAT; M-R-Cd300e CTT ACA CTG ACC GAT GGA TCA C; mTNF-FWD CGC TCT TCT GTC TAC TGA ACT TCG G, mTNF-REV AGA ACT GAT GAG AGG GAG GCC ATT T; mGapdh-FWD AAT GTG TCC GTC GTG GAT, mGapdh-REV CTC AGA TGC CTG CTT CAC; M-F-Marco CCT CCA GGG ACT TAC GGG T, M-R-Marco CCA GTG AGA CCT ATG TCA CCT; M-F-Slamf9 CAA AAC AAC ATT GCC ATC GTG A, M-R-Slamf9 GCT AAT ATG CAG GGA GTA GCT G; M-F-Cleac4a1 GAC TCG TCT TCA TGT ACC GTC T, M-R-Cleac4a1 AGC AAC AGA GAA TAA GAT TGC CA; M-F-Clec4a2 CCC CCA TTG GAC AAA GGG C, M-R-Clec4a2 GGT GCC AAG ATA CCC AAG TCT A

### RNA sequencing

Libraries were generated using an in-house multiplex RNA-seq method (Version 01/03/2024; Hudson Genomics Facility) adapted from^25^. Briefly 25 ng of total RNA from each sample was tagged with a 10 bp sample index and 10 bp unique molecular identifier (UMI) during initial poly(A) priming and pooled samples amplified using a template-switching oligo. After cDNA amplification Illumina P5 adaptors with distinct i5 indexes were added by tagmentation by Nextera transposase and PCR. The final libraries were pooled for subsequent processing and single-end sequencing was performed on a NextSeq 2000 run using a P2 100 cycle kit. For sequencing, the multiplex library was loaded at 900pM with ∼1% PhiX for on-board denaturation and clustering. Base calling was performed using Dragen BCLConvert (v4.2.7). Demultiplexing of raw sequencing files was performed by the sequencing facility.

### RNA-seq analysis

RNA-seq analysis was performed in R (v4.1.0, FASTQ file processing; v4.5.0, statistical analysis)^26^. The scPipe package (v1.14.0) ^27^ was employed to process the data. Read alignment was performed on R1 FASTQ files using the Rsubread package (v2.6.1)^28^. An index was built using the Ensembl *Mus musculus* GRCm39 primary assembly genome file and alignment was performed with default settings. Aligned reads were mapped to exons using the sc_exon_mapping function with the Ensembl *Mus musculus* GRCm39 v104 GFF3 genome annotation file. Reads mapping to exons in the resulting BAM files were associated with each individual sample using the sc_demultiplex function, taking the UMI into account, and an overall count for each gene for each sample was generated using the sc_gene_counting function (with UMI_cor = 1). Additional gene annotation was obtained using the biomaRt package (v2.66.2) ^29^ and a DGEList object was created with the counts and gene annotation using the edgeR package (v4.8.2) ^30^. A design matrix was constructed incorporating the treatment group and donor mouse (BMDMs from n=3 mice were used).

Lowly expressed genes were removed using the filterByExpr function and normalisation factors were calculated using the TMM method ^30^. Counts were transformed for differential gene expression analysis using the voom method ^31^ and a linear model was fit using the edgeR voomLmFit function. Differential gene expression tests were performed using the limma package (v3.66.0)^32^. Comparisons were made for each treatment group compared to non-treated controls using the contrasts.fit function and empirical Bayes moderated t-statistics were calculated using the eBayes function^33^. Differentially expressed genes were determined using a false discovery rate (FDR) adjusted p value < 0.05. Volcano plots were made using the log2 fold changes of all genes versus the –log10 p value across all samples as calculated during this step. The original demultiplexed R1 FASTQ files were deposited along with the UMI-adjusted counts in the NCBI Gene Expression Omnibus (GEO) with accession GSE327214.

### Recombinant TLR7 and TLR8 proteins

Preparation of recombinant TLR7 and TLR8 extracellular domain was described previously^9, 34^. *Macaca mulatta* TLR7 extracellular domain (N167Q, N399Q, N488Q, N799Q mutations, and residues 440-445 (SEVGFC) replaced by a thrombin-cleavage sequence (LVPRGS)) or *Homo sapiens* TLR8 extracellular domain fused to a C-terminal thrombin-cleavage sequence followed by a protein A tag were inserted into the pMT-BiP-V5-His vector (Thermo Fisher Scientific). For expression, stably transfected *Drosophila* S2 cells (Thermo Fisher Scientific) were cultured in EXPRESS FIVE SFM medium (Thermo Fisher Scientific) and induced by addition of 0.5-0.6 mM copper(II) sulfate to the culture medium. For TLR8 protein used for crystallization, kifunensine (1.7-1.9 mg/L), a mannosidase I inhibitor, was also added to the culture medium. Culture supernatant containing secreted proteins was harvested and subjected to affinity purification using IgG Sepharose 6 Fast Flow affinity resin (Cytiva). After affinity purification, proteins were treated by thrombin for Z-loop (TLR7) and tag cleavage. TLR8 proteins used for crystallization were additionally treated by endo Hf for glycan trimming. Proteins were purified using gel-filtration chromatography. Purified proteins were used for structural analysis and SPR experiments. TLR7 F507S and TLR8 F495S mutant proteins were described previously and prepared similarly^12^.

### Surface plasmon resonance (SPR)

Using a Biacore T200 (Cytiva), TLR7 proteins were immobilised onto a Series S sensor Chip CM5 (GE) by amine coupling to a level of between ∼12,000 RU for TLR7 and mutants and ∼12,000-14,000 RU for TLR8 and mutants, respectively, as per the manufacturer’s instructions. Briefly, at 25°C, with a flowrate of 10 µl/min, flow cells were activated with injection of 0.2 M EDC + 0.05 M NHS for 420 s. TLR7 proteins (300 µg/ml in 10 mM acetate pH 4) were coupled to the chip for 650 s at 2 μl/min), and TLR8 proteins (200 μg/ml in 10 mM Acetate pH 4) were coupled to the chip for 600 s at 2 μl/min. Unreacted NHS was blocked on all flow cells with an injection of 1 M ethanolamine-HCl pH 8.5 for 420 s. One flow cell was used as the reference, activated and blocked as described above. The immobilisation buffer was 20 mM HEPES, 150 mM NaCl, pH 7.5.

The SPR run was performed at 20°C with 10 mM Mes, 150 mM NaCl, pH 5.5 as the running buffer. All oligonucleotides were solubilised in running buffer, and concentration confirmed via the calculated extinction coefficient at A260 nm as well as 1D ^1^H NMR (for R848 only). A 6-point dilution series at the given concentration range was performed for all oligonucleotides. Samples were injected for 120 s with a dissociation time of 300 s at a flow rate of 40 µl/min. Surface regeneration was performed by 2 x 60 s injections of 2 M NaCl at 30 µl/min. Data were analysed using Biacore T200 Evaluation Software. Affinity constants were determined by fitting the data using affinity analysis with a standard 1:1 Langmuir binding model. R848 affinity binding or Circular Dichroism (CD) spectroscopy were used to confirm correct folding of all TLR7 and TLR8 mutant proteins, respectively.

### Isothermal Titration Calorimetry (ITC)

ITC experiments were carried out at 20°C in 10 mM Mes, 150 mM NaCl pH 5.5 using a Malvern MicroCal PEAQ-ITC (Malvern Instruments). Protein was buffer exchanged using a Zeba^TM^ Spin desalting column (7K MWCO, ThermoFisher) into the ITC buffer, and oligonucleotides were prepared from powder into the same buffer stock and quantified using the calculated extinction coefficient at A260 nm.

The titration sequence included a single 0.4 μL injection, followed by 18 injections, 2 μL each with a 120 s interval between injections. The titration conditions were 350 μM oligonucleotide into 30 μM TLR7. Data were analysed using the MicroCal PEAQ-ITC Analysis software.

### Cryo-EM analysis, data processing and model building

Recombinant TLR7 extracellular domain (0.2 mg/mL) was mixed with GUC-v16 (0.17 mg/mL) (WuXi AppTec) using a buffer containing 20 mM sodium citrate, pH 5.0 and 150 mM NaCl. A 3-μl aliquot was applied onto a glow-discharged QUANTIFOIL® R 1.2/1.3 on Cu 300 mesh grid + 2 nm C Holey Carbon Films with ∼ 2 nm continuous carbon on top (QUANTIFOIL, #C2-C14nCu30-50). The grids were blotted for 2.0 s in 100% humidity at 6°C and plunged into liquid ethane using a VitrobotMkIV (Thermo Fisher Scientific). Cryo-EM movies were recorded by using a Titan Krios G4 microscope (Thermo Fisher Scientific) running at 300 kV and equipped with a Gatan Quantum-LS Energy Filter (GIF) and a Gatan K3 camera in the electron counting mode at the Cryo-EM facility in the University of Tokyo (Tokyo, Japan). Imaging was performed at a nominal magnification of 105,000×, corresponding to a calibrated pixel size of 0.83 Å/pixel. Each movie was recorded for 1.6 s and subdivide into 48 frames with an accumulated exposure of about 48 e^−^/Å^2^. EPU software (Thermo Fisher Scientific) operated at the fast acquisition mode was used for data collection.

For processing of the cryo-EM data, 3,602 raw movies were imported in RELION v4.0.1^35, 36^, and were motion-corrected using MotionCor2 in RELION’s own implementation^37^. The CTF parameters were determined using the CTFFIND4 program^38^. Template-free auto-picking using the Laplacian-of-Gaussian filter was performed for particle picking. Picked particles were extracted and divided into 20 subsets. A single round of 2D classification was performed to exclude junk particles. Multiple rounds of 3D classification, 3D auto-refine, Post-processing, Bayesian polishing and CTF refinement were performed to select the final particle stacks containing 169,275 particles for the final 3D auto-refine and Post-processing. Local resolutions of the final 3D map were estimated using RELION’s own implementation. The detailed cryo-EM data processing flowchart is shown in Extended Data Figure 5. For model building, the TLR7 structure (PDB 9LUV) was used as the initial model, which was fitted into the cryo-EM 3D map of TLR7/GUC-v16 complex using the ‘fit in map’ tool in Chimera software^39^. Restraints (CIF file) for GUC-v16 was generated using the JLigand software implemented in the CCP4 program suite ^40^. The iterative cycles of manual model building in COOT ^41^ and real-space refinement in Phenix software ^42, 43^ were continued until the structures converged with reasonable geometric parameters and map-to-model fit. Both the unsharpened and the B-factor sharpened 3D maps were referred during manual refinement using COOT, and the unsharpened 3D map was used in real-space refinement in Phenix. The 3D maps and atomic coordinates have been deposited in the Electron Microscopy Data Bank (EMD-80227) and Protein Data Bank (PDB: 25NO), respectively. Statistics for data collection and structural refinement are summarized in Table S4. Structure representations were generated using Chimera^39^, ChimeraX^44^ and COOT ^41^.

### Crystallization, data collection and structure determination

Crystallization experiment was conducted using the sitting-drop vapor-diffusion methods at 20°C. Droplet setting was performed using mosquito crystal (TTP Labtech). For co-crystallization of TLR8 and GUC-v16, protein solution containing 8 mg/mL TLR8 extracellular domain with 1.75 mg/mL GUC-v16 (WuXi AppTec) in a buffer containing 10 mM Tris-HCl pH7.5 and 0.15 M NaCl was used. 1.0-μL aliquots of protein solution and 1.0-μL aliquots of reservoir solution containing 0.1 M Tris-HCl, pH8.2, 0.2 M NaCl and 11.5%-17% PEG8000 (Hampton Research) were mixed to making the droplets. For co-crystallization of TLR8, uridine and mG*dC*dC (* is PS), protein solution containing 7.5 mg/mL TLR8 extracellular domain with 0.41 mM uridine and 0.41 mg/mL mG*dC*dC (WuXi AppTec) in a buffer containing 10 mM Tris-HCl pH7.5 and 0.15 M NaCl was used. 0.8-μL aliquots of protein solution and 0.8-μL aliquots of reservoir solution containing 0.1 M sodium citrate, pH5.0, 0.2 M potassium formate and 12% PEG3350 (Hampton Research) were mixed to making the droplets. Crystals were soaked into cryoprotectant solutions containing 15% glycerol and were mounted with nylon loops and flash cooled using liquid nitrogen. X-ray diffraction experiments were conducted by using the automatic data collection system ZOO ^45^ at SPring-8 BL32XU beamline under the cryogenic condition of 100 K with a wavelength of 1.0000 Å. Diffraction data were automatically processed by using the XDS ^46^ and KAMO program ^47^. For the TLR8/GUC-v16 complex, data collected from multiple crystals were clustered and merged with equivalent unit cell parameters by using the KAMO program ^47^ and a merged dataset was selected. For the TLR8/U/mG*dC*dC complex, diffraction data were collected from a single crystal. The phases were determined by molecular replacement method using the MOLREP program ^48^ implemented in the CCP4 program suite. TLR8/uridine complex structure (PDB 4R0A) chain A was used as the search model. Restraints (CIF file) for mGdCdC was generated using the JLigand software implemented in the CCP4 program suite ^40^. The structure was refined with stepwise cycles of manual model building using the COOT ^41^ and REFMAC5 ^49^ programs. Atomic coordinates and structure factors of TLR8/GUC-v16 complex and TLR8/U/mG*dC*dC have been deposited in the Protein Data Bank (PDB: 25NM and PDB: 25NN, respectively). The statistics of the data collection and refinement are summarized in Table S2. Structure and electron density representations were generated using ChimeraX ^44^ and COOT ^41^.

### Homology modelling

The cryo-EM structure of mmTLR7 in complex with GUC-v16 reported in this study was used to create the homology model of wild-type mouse TLR7 inactive dimer in complex with GUC-v16 using MODELLER (version 10.4)^50^.

### Mouse studies

All the animal experiments complied with relevant local ethical regulations.

### Aldara-driven skin inflammation model

These experiments were approved in advance by an Animal Ethics Committee at Monash Medical Centre (MMCB/2023/19) and were carried out in accordance with “Australian Code of Practice for the Care and Use of Animals for Scientific Purposes. Eight week old C57Bl/6J female mice were used in these experiments and were housed in SPF (Specific Pathogen Free) conditions. Mice were anaesthetized for 1-2 minutes with a 26aporizer machine (oxygen flow rate of 1-4 litres/min) with fresh oxygen and 5% isoflurane. Upon induction of anesthesia, back hair was removed using sensitive hair removal cream (Nair). A small drop of Nair cream was applied to mouse’s upper back using a cotton bud and smeared to cover a 2 cm^2^ patch. The Nair cream was allowed to sit for ∼one minute before being wiped off in a nose-tail motion using a sterile cotton bud to remove hair from the back skin. Thereafter, mice were treated topically to the hairless back skin with 100 ml of GUC-v16 (98.7% purity) formulated as a 2.5% w/w cream composed of a mixture of excipients, including water, propylene glycol, Polawax™ NF, Caproyl 90, stearyl alcohol, Laurogylcol FCC, and Tween 80, and allowed to absorb for 1 min. 50 mg of Aldara cream (containing imiquimod, 5% w/v) was subsequently applied to the back skin using a cotton bud. One cohort of mice was treated with Vaseline cream as vehicle control for the Aldara cream. These treatments were repeated daily for 4 consecutive days. Mice were scored daily for back erythema (redness) and skin scaling, as previously reported ^51^. Spleens and skin samples were collected at the end of the experiment (day 5).

### RNA purification from skin biopsies

Skin samples were cut into small pieces and suspended in 300 μl of lysis buffer (PureLink RNA Mini Kit – Thermo Fisher Scientific), prior to homogenisation on ice in 5-15 sec bursts for 2-5 min using IKA T10 homogeniser (T10 basic ULTRA-TURRAX). The homogenates were subsequently centrifuged at 12,000 x g for 5 minutes and the supernatants were transferred to a clean Rnase-free tubes and total RNA extracted with the PureLink RNA Mini Kit (Thermo Fisher Scientific).

### Systemic R848 challenge

8-week old C57BL/6NCrl mice (used under Australian National University animal ethics, reference A2023/309) were injected intravenously with 200 μg of GUC-v16 conjugated with *in vivo*-jetPEI® (Polyplus #101000030) in 5% Glucose, 1 h prior to intraperitoneal injection with 25 μg R848 VacciGrade (InvivoGen #vac-r848). Two hours after injection of R848, mice were sacrificed for spleen collection. The spleen was processed to a single-cell suspension and following lysis of red blood cells, total RNA was purified from 5×10^6^ total splenocytes using the Isolate II RNA mini-kit (Meridian Biosciences # BIO-52072) according to the manufacturer’s instructions. RNA was transcribed into cDNA using SuperScript IV Reverse Transcriptase (Thermo Fisher Scientific #18090010) according to the manufacturer’s instructions. RT-qPCR analyses were carried out with the Power SYBR Green Master Mix (Thermo Fisher Scientific) on an Applied Biosystems 7900 machine (Thermo Fisher Scientific). Each PCR was performed in technical triplicates with mouse *Gapdh* used as the reference gene. Relative gene expression was calculated using the 2^-ΔΔCt^ method.

### Intratracheal R848 challenge

Eight-ten week-old female C57BL/6J mice were purchased from Janvier Labs (France) and housed under standard conditions with ad libitum access to dry food and water. All experimental procedures complied with EU Directive 2010/63/EU on the protection of animals used for scientific purposes and were approved by the local ethics committee (CEBEA; approval number 51 Gos CMMI). Mice received 2.5 µg of naked GUC-v16 in water via endotracheal administration under isoflurane anesthesia (4% for induction and 2% for maintenance) using a dedicated endotracheal device (Fine Mist Sprayer for mice, Aptar, France) following the manufacturer’s instructions. One hour later, mice were administered 50 µg of R848 VacciGrade (InvivoGen, #vac-r848) in PBS via endotracheal delivery. Seven hours after R848 administration, animals were euthanized and bronchoalveolar lavage and blood were collected. For BAL collection, two sequential lavages with 800 µL of PBS were carried out. Cell density in BAL fluids was assessed using a Thoma counting chamber, and differential cell counts were performed on cytospin preparations (≥200 cells per sample) stained with RAL 555 May–Grünwald–Giemsa (RAL, #720-0351). Cytokine levels in BAL and plasma were quantified using an MSD (Meso Scale Discovery) multiplex assays for mouse IFN-β, IL-10, IL-6, IP-10, KC/GRO (CXCL1), MIP-2 (CXCL2) and TNF-α (U-PLEX Custom Biomarker Group 1 (mouse) Assays, #K15069M-1, batch 574581) on a MESO QuickPlex SQ 120MM instrument carried, out by the CER group (Belgium) under investigation study 1030_2026-INH-02. Data were processed using the DISCOVERY WORKBENCH Desktop Analysis Software. Unlike plasma samples, which required no additional treatment, BAL samples were supplemented with 1% Blocker A (MSD, #R93AA-2), to ensure optimal assay performance and to prevent analyte loss due to adsorption to labware.

## Statistical analyses

Statistical analyses were carried out using Prism 10 (GraphPad Software Inc.). One-way and two-way analyses of variance (ANOVA) with uncorrected Fisher’s LSD were used when comparing groups of conditions, while unpaired two-tailed t-tests were used when comparing selected pairs of conditions. “ns” is non-significant.

## Acknowledgements

We thank K. Sakaniwa for the purified TLR8 protein; M. Sweet for the RAW-ELAM cells; E. Bartok for TLR7-deficient THP-1 cells^21^; T. Wilson for RNA sequencing and help with analyses; J. Lamb for technical assistance with mouse experiments; the CER group for performing the cytokine analyses in plasma and BALs using the MSD assay; M. Kikkawa and Y. Sakamaki for managing and supporting the Graduate School of Medicine cryo-EM facility at the University of Tokyo. We thank the Beamline staff members at SPring-8 for their assistance with data collection. Especially, we thank K. Hirata, Y. Kawano and H. Matsuura for automated data collection at SPring-8 BL32XU. We acknowledge use of the Sydney Analytical Core Research Facility (University of Sydney) for access to SPR and ITC infrastructure.

This work was supported by the Australian National Health and Medical Research Council Project Grant (2020565 to M.P.G., J.I.E and B.C.); mRNA Victoria Research Acceleration Fund, the Victorian Government’s Operational Infrastructure Support Program and COVID-19 Treatments Medical Research Fund; Deutsche Forschungsgemeinschaft (DFG) grants CRC237 369799452/B21 (M.A.L.K.), CRC237 369799452/A11 (M.A.L.K.), CRC369 501752319/C06 (M.A.L.K.), and CRC237 369799452/A06 (C.W.); the German Federal Ministry of Research, Technology and Space (BMFTR) grant 01GL2405B (M.A.L.K.) as part of the German Center for Child and Adolescent Health (DZKJ); Grant-in-Aid from the Japanese Ministry of Education, Culture, Sports, Science, and Technology (Grant Nos. 24K09349 and 26H01561 to Z.Z., 25K09524 to A.F., 25K02214 and 25H01843 to U.O., 22H05184, 23H00366 and 25K22517 to T.S.); CREST, JST (Grant No. JPMJCR21E4 to T.S.); the Francis Crick Institute (CC2228), which receives its core funding from Cancer Research UK, the UK Medical Research Council, and the Wellcome Trust; and Noxopharm Limited. Cryo-EM analyses were supported by the Basis for Supporting Innovative Drug Discovery and Life Science Research (BINDS) from the Japan Agency of Medical Research and Development (AMED) (Grand No. JP21am0101115; support No. 1570, 1846, 1848). X-ray diffraction experiments were supported by the BINDS from the AMED (Grand No. JP23ama121001; support No. 5218). This research project was undertaken with the assistance of resources and services from the National Computational Infrastructure and the Pawsey Supercomputing Research Centre, which are supported by the Australian Government and the Government of Western Australia accessed through the National Merit Allocation and ANU Merit Allocation schemes. This work was also supported by the Region Wallonne Win4Company convention 8666. The LSR FORTESSA Flow Cytometer was supported by DFG grant 446167311. This work was supported by Monash eResearch, a node of the ARDC Nectar Research Cloud.

## Author contributions

Conceptualization: M.P.G, S.S., O.F.L., D.S.W., M.S., Z.Z., T.S., J.I.E., C.W., M.A.L.K., P.P., R.R.; Investigation: S.S., Z.Z., C.W., A.F., L.W.W., E.R., N.B., P.P., A.L., L.C., R.G., P.M.S., W.S.N.J., J.I.E., A.L.M., C.C., R.G., M.S., D.S.W., J.B., A.H., A.S.A., U.O., R.J., B.C., M.A.L.K., O.F.L., M.P.G.; Resources: M.W., K.A.L., M.A.B., U.O., O.F.L., D.S.W., M.S., C.R., J.I.E, M.A.L.K., R.R., B.C., T.S., M.P.G.; Data Curation: A.L.M., Z.Z.; Writing – Original Draft: M.P.G., S.S., D.S.W., Z.Z., T.S.; Writing – Review & Editing: M.P.G., S.S., D.S.W., L.W.W, L.C., R.G., B.C., A.L.M., M.S., P.P., C.R., M.A.L.K., C.W., J.I.E., R.J., K.A.L.; Supervision: M.P.G., T.S., O.F.L., J.I.E., M.A.L.K., R.G.; Project administration: S.S., Z.Z., U.O., C.W., D.S.W, M.S., J.I.E., R.G., O.F.L., M.A.L.K., T.S. and M.P.G.; Funding acquisition: C.W., C.R., R.R., M.A.L.K., O.F.L., J.I.E., T.S., M.P.G.

## Data and materials availability

RNA sequencing data has been deposited in the NCBI Gene Expression Omnibus (GEO) with accession GSE327214. The coordinates and structure factors have been deposited at the Protein Data Bank (PDB) under the following accession codes: TLR8/GUC-v16 complex (25NM) and TLR8/U/mGdCdC (25NN). The cryo-EM map and coordinates for the TLR7/GUC-v16 complex have been deposited at the Electron Microscopy Data Bank and PDB under the accession codes: EMD-80277 and 25NO, respectively.

**Extended Data Figure 1.**
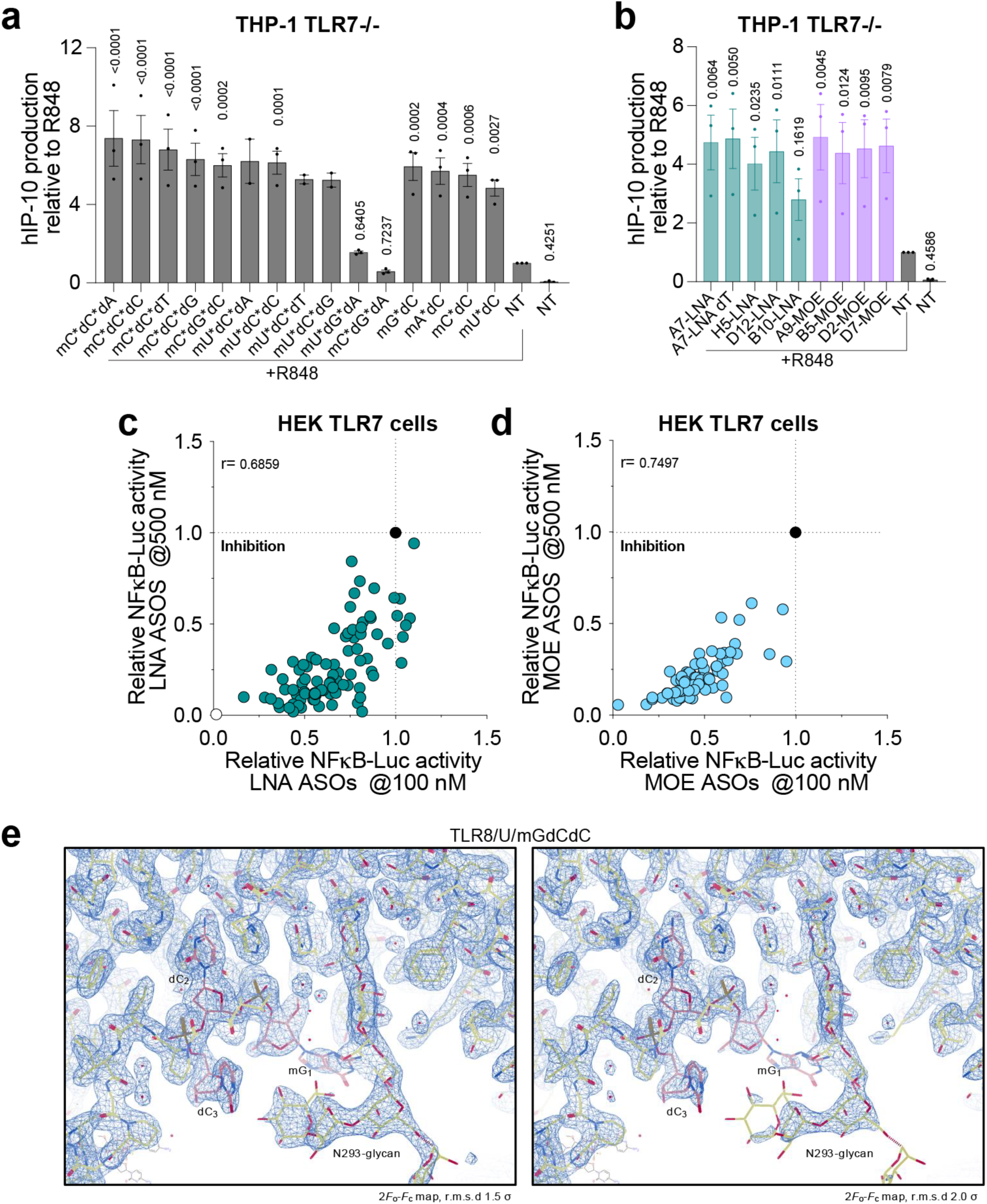
TLR7 and 8 modulation by oligos. (**a** and **b**) Monocytic THP-1 TLR7KO cells were incubated overnight with 5 μM (**a**) and 500 nM (**b**) of indicated 2-mers, 3-mers and 20-mers oligos and stimulated with 1 μg/ml R848 for 8 h prior to IP-10 ELISA analysis. Data are mean of n=3 independent experiments. IP-10 levels were normalised to the R848 only condition (± s.e.m. and two-way ANOVA with uncorrected Fisher’s LSD tests shown compared to the R848 only condition; a:*p<0.0001; b: p=0.0056*). (**c** and **d**) HEK TLR7 cells were pre-treated ∼30 min with 100 nM and 500 nM of indicated oligos and stimulated with 1 μg/ml R848 overnight prior to luciferase assay. Data were background-corrected using non-treated (NT) condition and are shown as relative expression to R848 only. Data are averaged from 2 or 3 biological replicate for each screen, and the screens at the different oligo concentrations were conducted on independent days (*r* values are provided on each graph; c,d:*<0.0001*). (**e**) Electron density map (2*F*_o_-*F*_c_ map) of the TLR8/U/mGdCdC complex around site 2 at contour levels of r.m.s.d 1.5 σ (left panel) and r.m.s.d 2.0 σ (right panel).

**Extended Data Figure 2.**
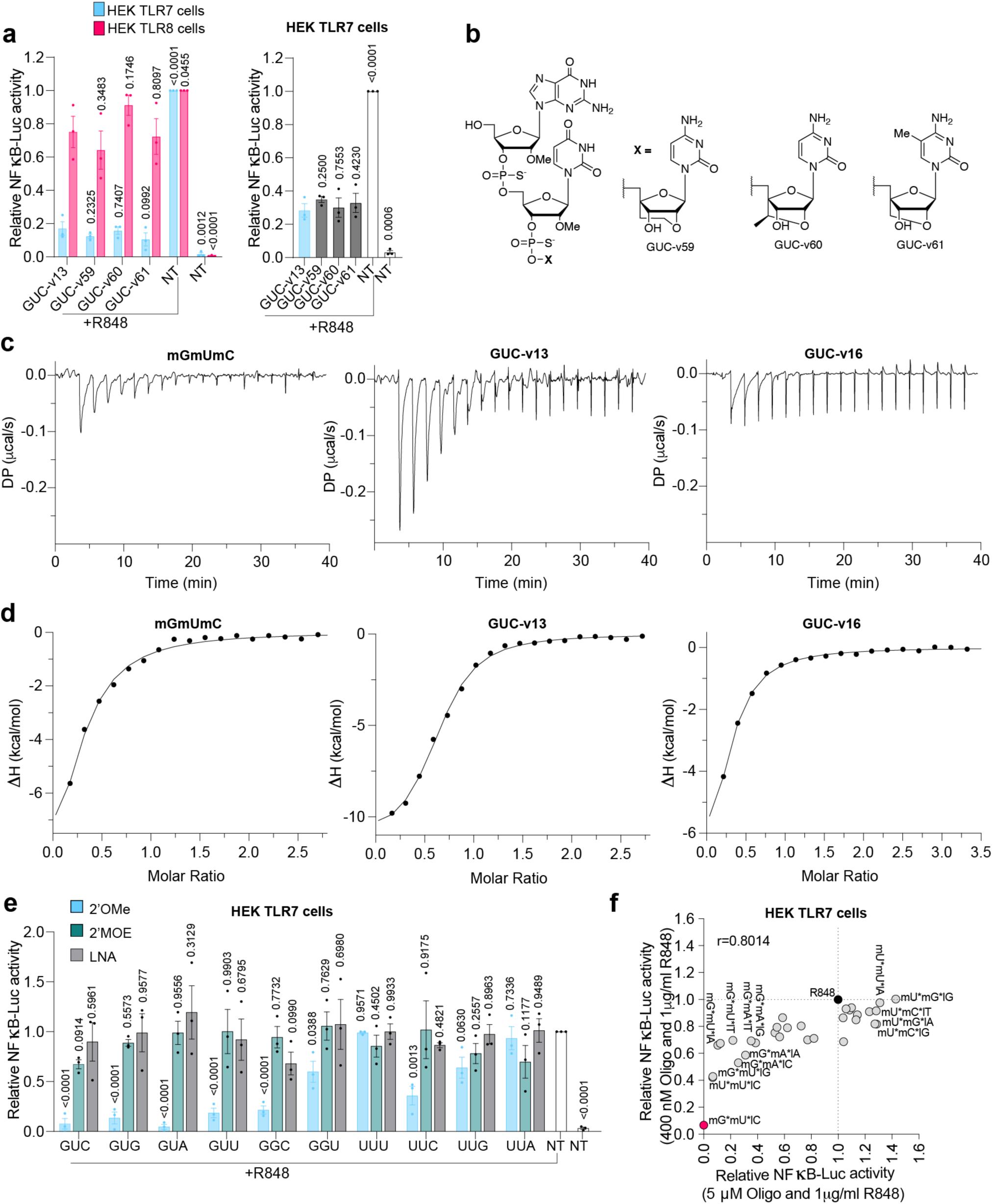
TLR7/8 modulation by modified 3-mer oligos. (**a** – left panel) HEK-TLR7 and HEK-TLR8 cells were pre-treated ∼30 min with 125 nM (TLR7) and 5 μM (TLR8) of the indicated oligos prior to overnight stimulation with 1 μg/ml of R848 followed by luciferase assay. Data are mean of n=3 independent experiments. Data were background-corrected using the non-treated (NT) condition and are shown as relative expression to R848 only (± s.e.m. and one-way ANOVA with uncorrected Fisher’s LSD tests shown compared to GUC-v13+R848 condition; *P<0.0001*). (**a**-right panel and **e**) HEK-TLR7 cells were pre-treated ∼30 min with 50 nM (**a**-right panel) and 2 μM (**e**) of the indicated oligos prior to overnight stimulation with 1 μg/ml of R848 followed by luciferase assay. Data are mean of n=3 independent experiments. Data were background-corrected using the non-treated (NT) condition and are shown as relative expression to R848 only (± s.e.m. and one-way ANOVA with uncorrected Fisher’s LSD tests shown compared to GUC-v13+R848 [a right panel] and R848 only [**e**] condition; **a**-right panel and **e**:*P<0.0001*). (**c**-**d**) isothermal titration calorimetry of the binding between recombinant mmTLR7 with indicated 3-mer oligo. K_D_ were _m_G_m_U_m_C: 6±1 μM; GUC-v13: 2±0.2 μM; GUC-v16: 4±0.2 μM. (**c**-**d**) Data are from one ITC analysis with the same batch of mmTLR7 protein. (**f**) HEK TLR7 cells were pre-treated ∼30 min with 400 nM and 5 μM of indicated oligos and stimulated with 1 μg/ml R848 overnight prior to luciferase assay. Data were background-corrected using non-treated (NT) condition and are shown as relative expression to R848 only. Data are averaged from 2 or 3 biological replicate for each screen, and the screens at the different oligo concentrations were conducted on independent days (*r* value is provided on the graph, and correlation *P* value is <0.0001).

**Extended Data Figure 3.**
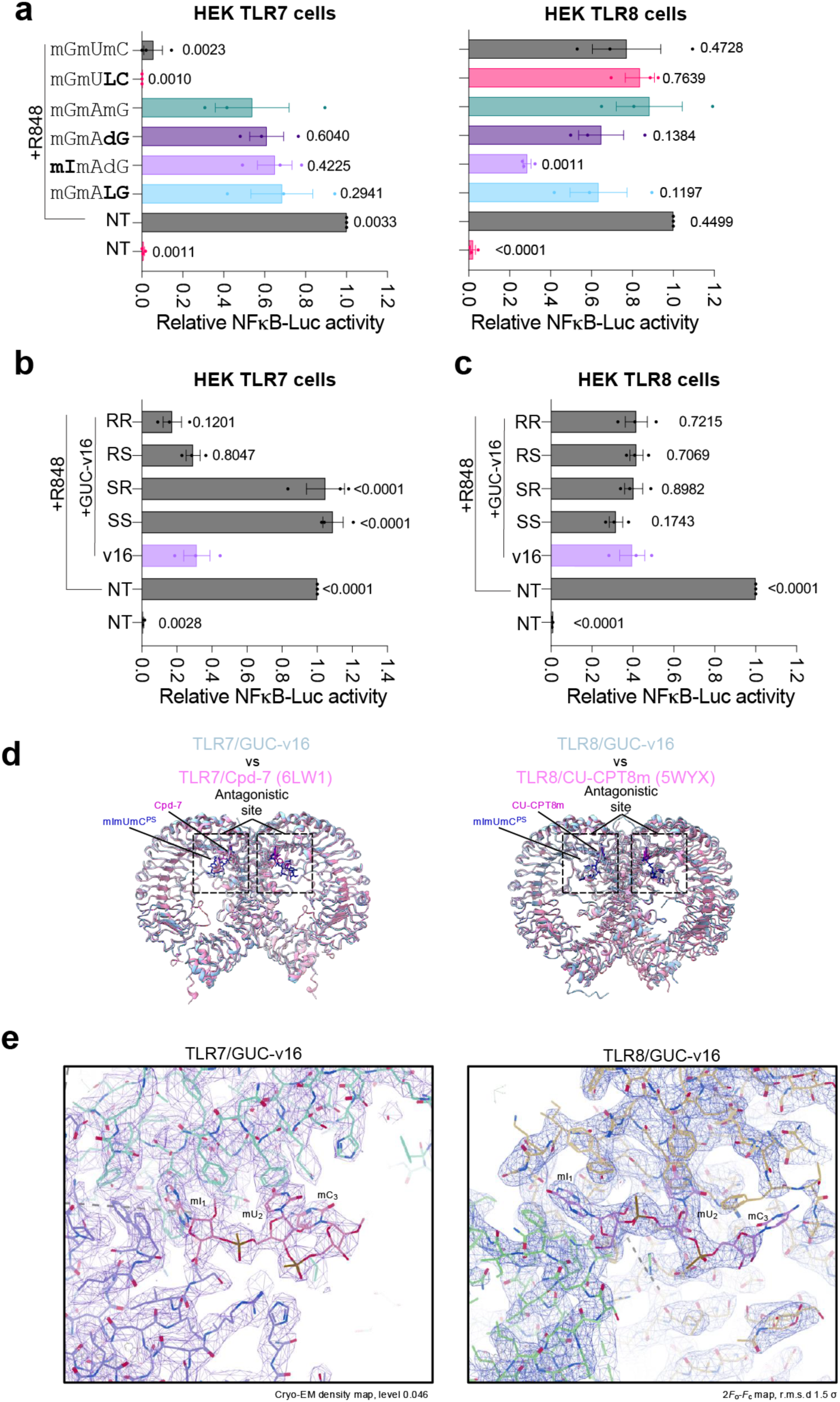
Inosine modulation of TLR7/8 sensing: **(a**, **b** and **c)** HEK-TLR7 and HEK-TLR8 cells were pre-treated ∼30 min with 200 nM (**b**), 1 μM (**a**) and 5 μM (**c**) of the indicated oligos prior to overnight stimulation with 1 μg/ml of R848 followed by luciferase assay. Data are mean of n=3 independent experiments. Data were background-corrected using the non-treated (NT) condition and are shown as relative expression to R848 only (± s.e.m. and one-way ANOVA with uncorrected Fisher’s LSD tests shown compared to GAG+R848 (**a**), GUC-v16+R848 (**b** and **c**) condition; **a** TLR7:*P<0.0001*; TLR8:*P<0.0001*; **b**:*P<0.0001*; **c**:*P<0.0001*). (**d**) Structural comparison of the TLR7/GUC-v16 and TLR7/Cpd-7 complex (PDB: 6LW1) structures (left panel) and the TLR8/GUC-v16 and TLR8/CU-CPT8m (5WYX) complex structures (right panel). (**e**) Cryo-EM density map (unsharpened map, contour level 0.046) of the TLR7/GUC-v16 complex (left panel) and electron density map (2*F*_o_-*F*_c_ map, contour level r.m.s.d 1.5 σ) of the TLR8/GUC-v16 complex (right panel) around the antagonistic sites.

**Extended Data Figure 4.**
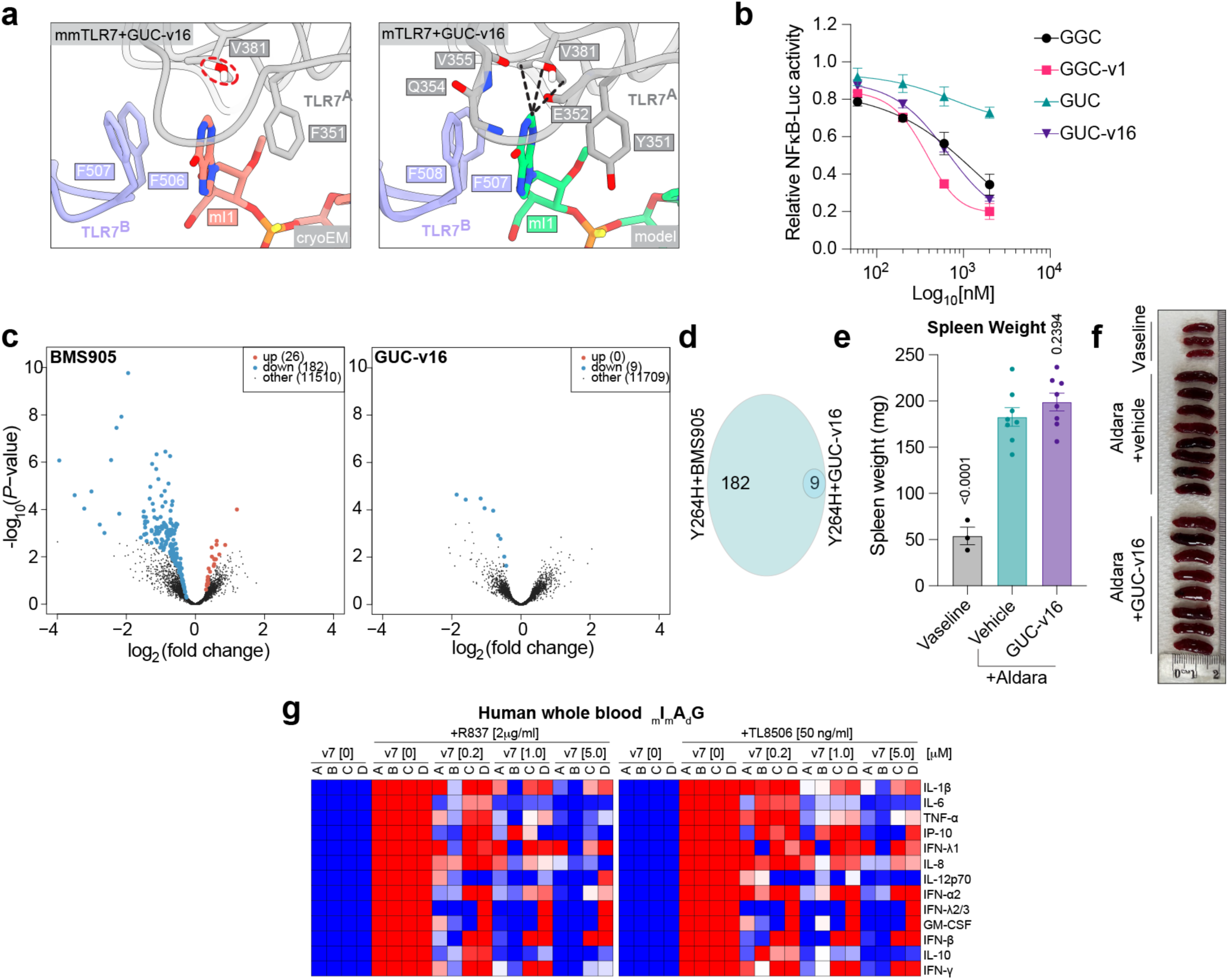
GUC-v16 modulated basal TLR7 sensing. (**a**) Close-up views of the antagonist binding site in the solved structure of mmTLR7 in complex with GUC-v16 (left) and the homology model structure of mouse TLR7 in complex with GUC-v16 (right). Residues shown in stick representations are selected residues form either hydrophobic interactions or hydrogen bonds with the first nucleoside base of 3-mer. Water molecule within dashed red circle is the water revealed in the cryo-EM structure of mmTLR7/GUC-v16 complex. Black dashed lines represent the newly formed hydrophobic interactions between the inosine base and V381 side-chain. (**b**) RAW-ELAM macrophages were pre-treated 1 h with different doses (2000 nM, 600 nM, 200 nM, and 60 nM) of the indicated oligos prior to overnight stimulation with 0.125 μg R848 followed by luciferase assay. Data are mean of n=3 independent experiments. Data were background-corrected using the non-treated (NT) condition and are shown as relative expression to R848 only (± s.e.m. and one-way ANOVA with uncorrected Fisher’s LSD tests shown compared to R848 only condition). (**c** and **d**) Bone-marrow-derived macrophages (BMDMs) from *Tlr7^Y264H^* mice were stimulated for 24 h with 1 μM of BMS905 and 5 μM of GUC-v16 oligo prior to RNA purification for RNA-sequencing. (**c**) Volcano plot of the genes significantly impacted by BMS905 and GUC-v16 treatments compared to non-treated condition (blue are down-regulated and red are upregulated) (**d**) Venn diagram of significantly different genes (the 9 common genes are: *Slamf9*, Nfkbiz*, Fpr1*, Clec4a1*, Slc13a3*, Serp1, Clec4a2*, Glrx, Rac2;* *= validated by RT-qPCR). (**e-f**) WT C57/BL6 mice were treated with Aldara cream directly following, or not, application of 100 μl of highly pure 2.5% GUCv-16 cream oligonucleotide (>99.4%). After four days, mice were humanely euthanised and spleen collected, weighed (e) and photographed with a ruler (cm) (**f**). (**e**) Mean of n = 3-8 mice/group is shown (± s.e.m. and one-way ANOVA with uncorrected Fisher’s LSD tests shown compared to Aldara group). (g) Whole blood from 4 healthy controls was pre-treated with indicated concentration of _m_I_m_A_d_G concentration for 30 minutes prior to stimulation with 2 μg/ml of R837 or 50 ng/ml TL8-506 for 24 h prior to cytokine bead analyses. For each control, cytokine levels were averaged for each technical replicate (6/sample), background corrected to unstimulated samples only, and reported to cytokine levels from stimulation only controls. Heat maps were generated using scale from 0-1 (see methods), were 1 is the cytokine level of the agonist only control. Data shown are from a minimum of two independent experiments – conducted on two independent days.

**Extended Data Figure 5:**
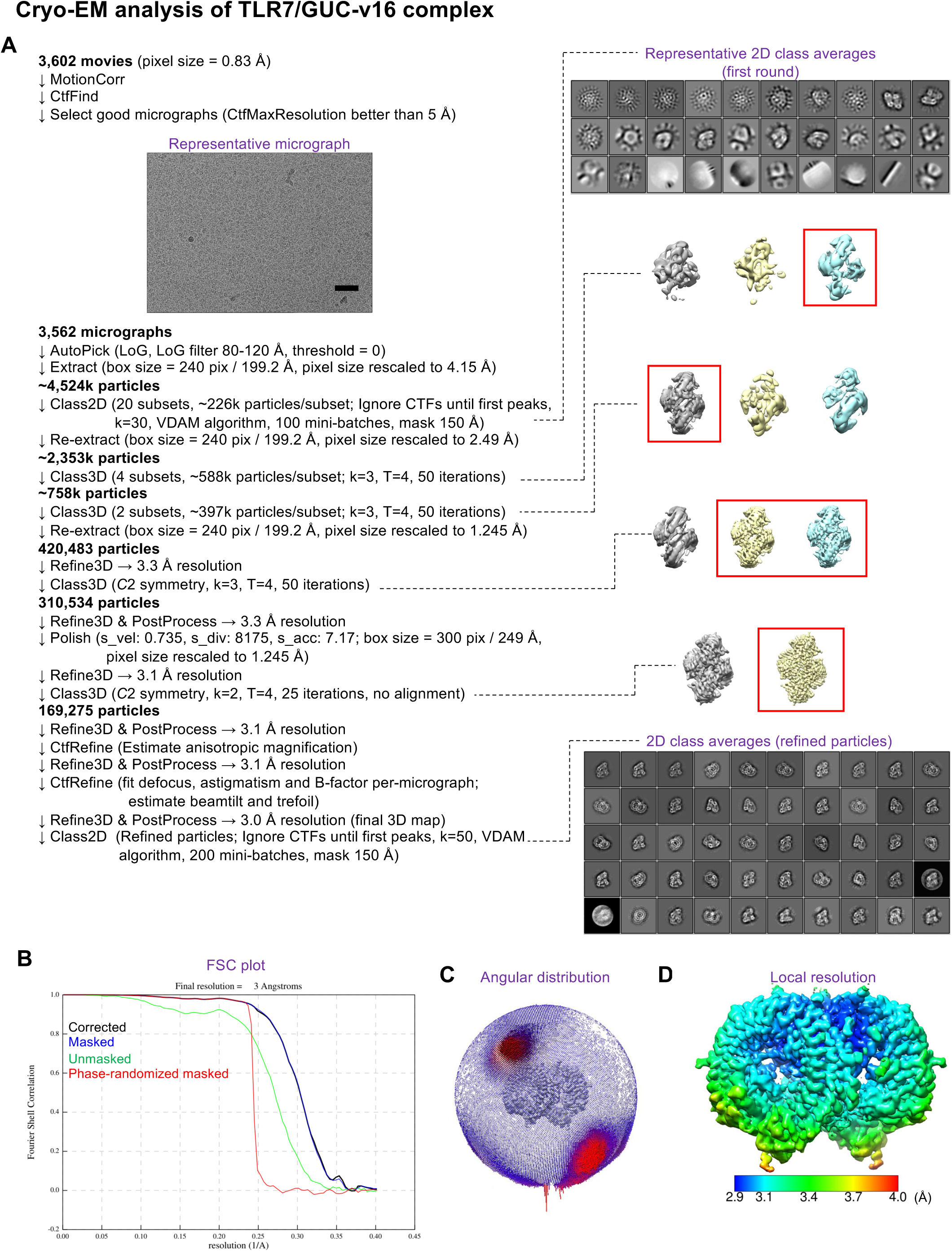
Cryo-EM analysis of TLR7/GUC-v16 complex. (a) Data processing workflow of cryo-EM analysis of the TLR7/GUC-v16 complex. Representative motion-corrected micrographs (out of 3,602 total micrographs), representative 2D class averages and 3D classes are shown. Fourier shell correlation (FSC) plot (a) of the final 3D reconstruction (resolution cut-off at FSC = 0.143), angular distribution (b) of the final refined particles and local resolution (c) of the final 3D map are shown.

**Supplementary Table S1:**
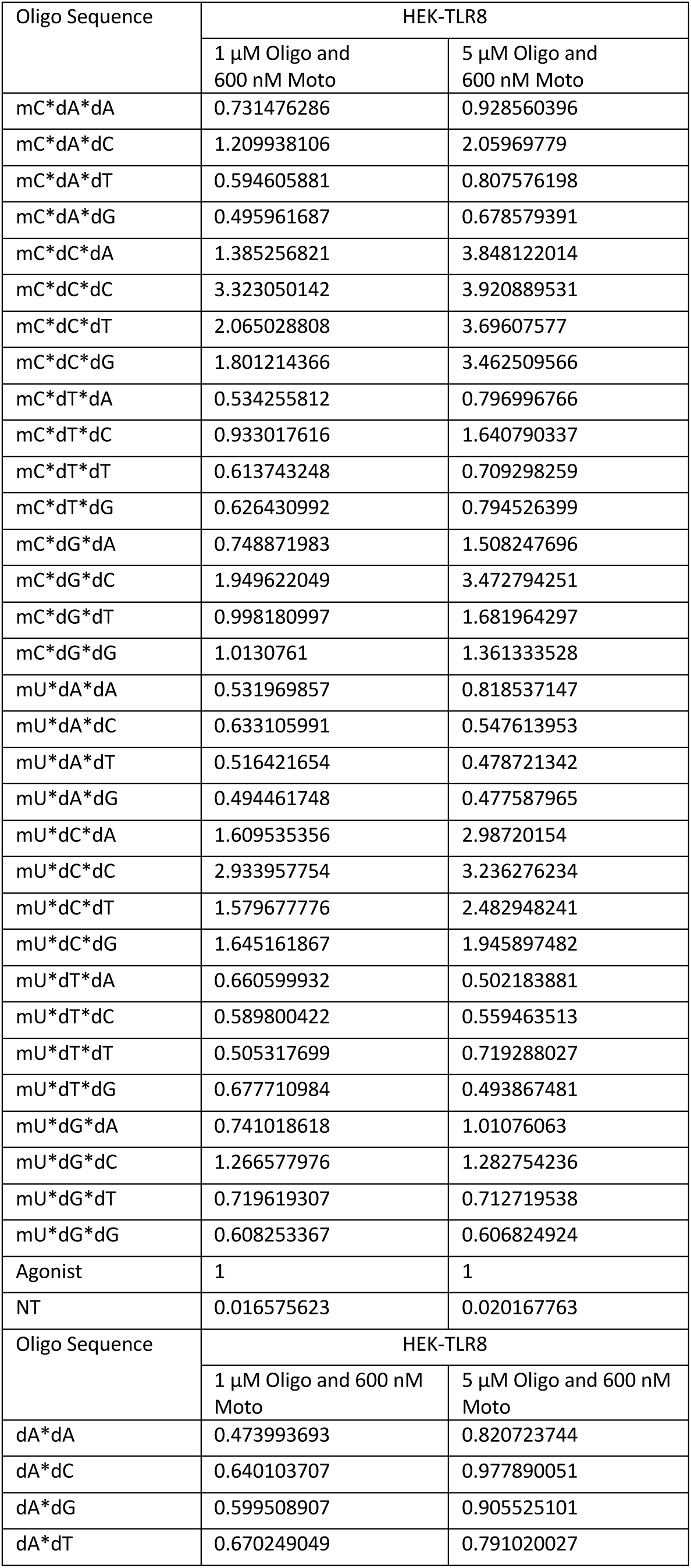
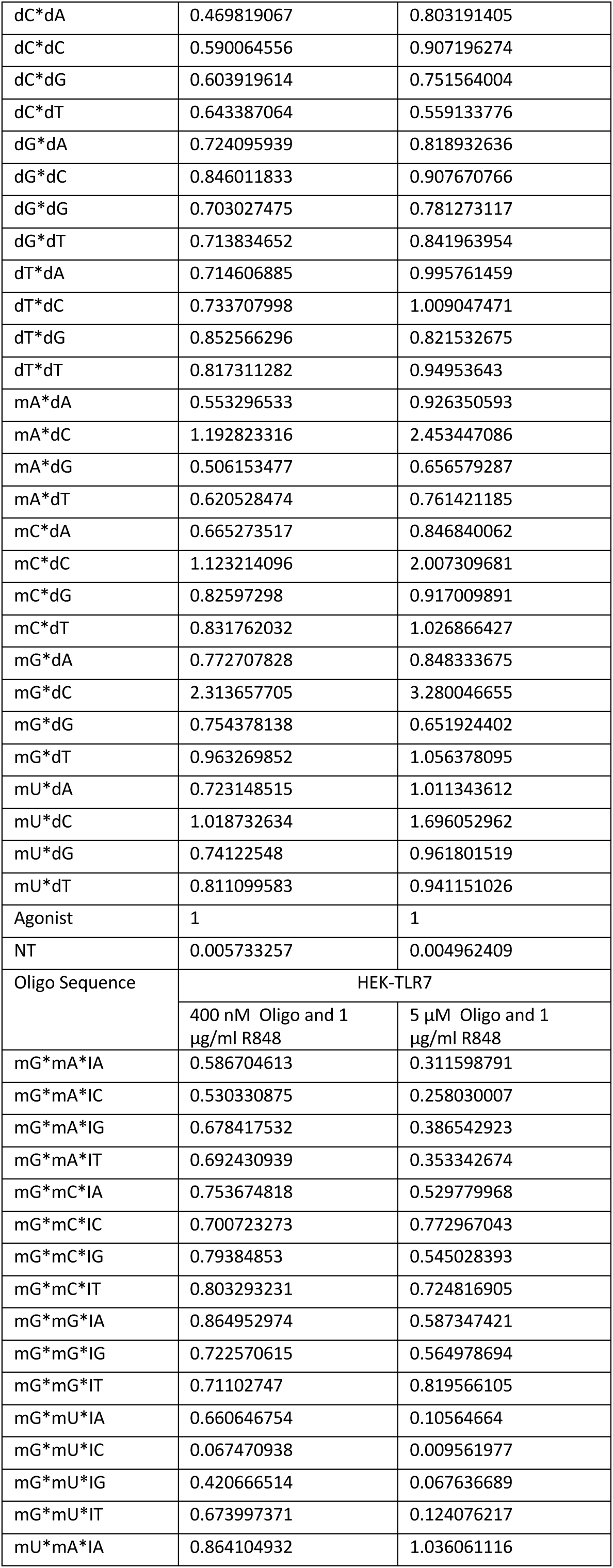
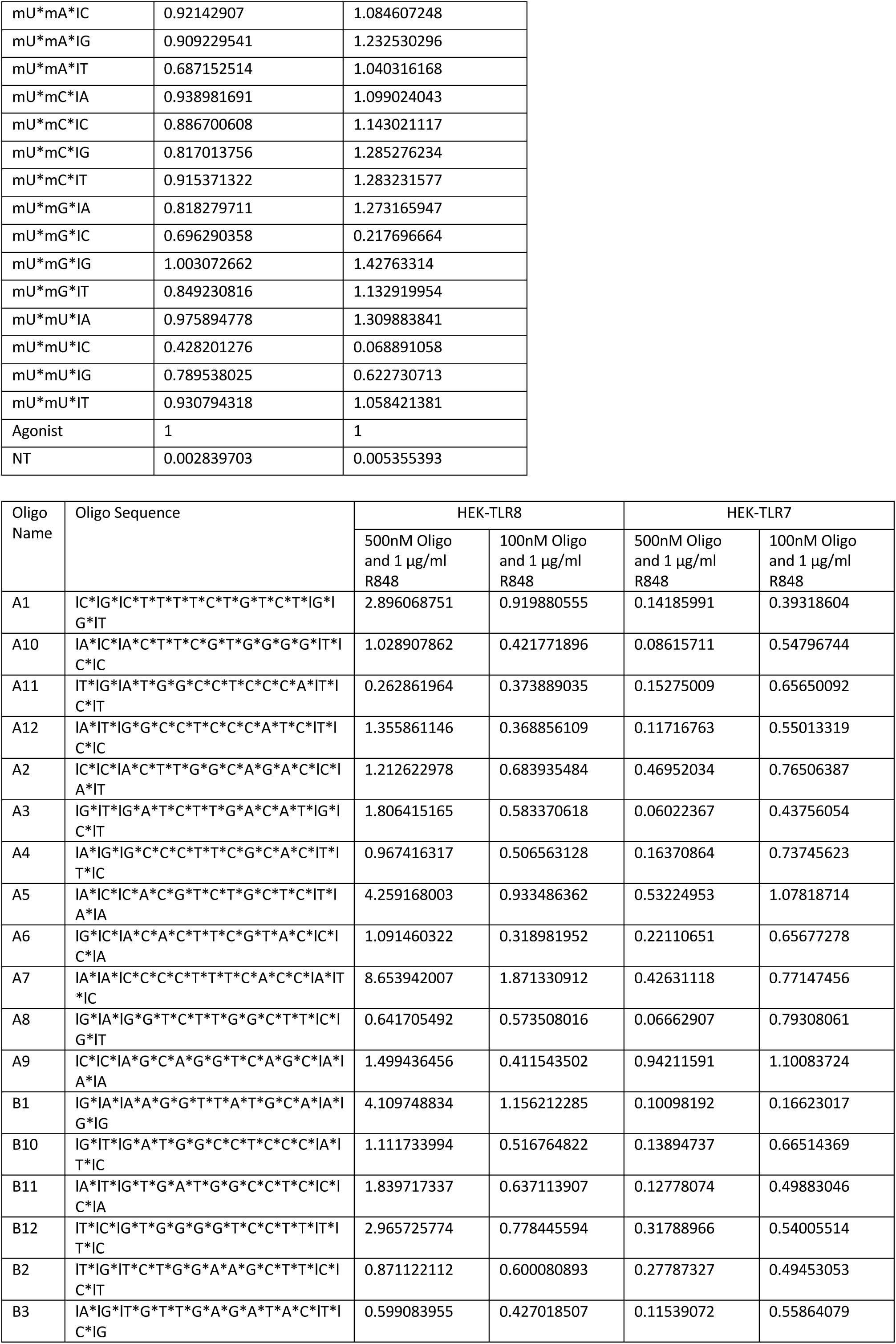
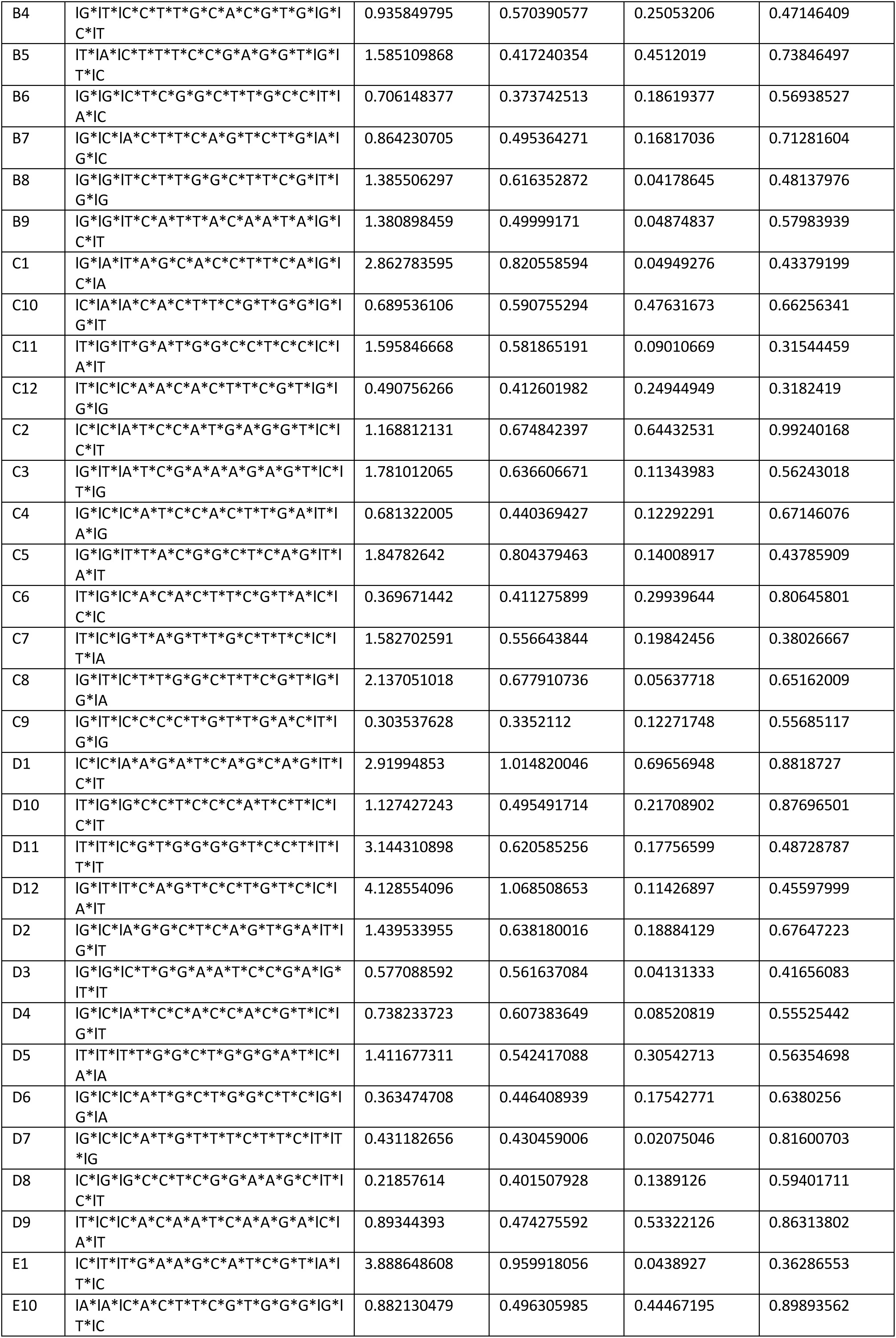
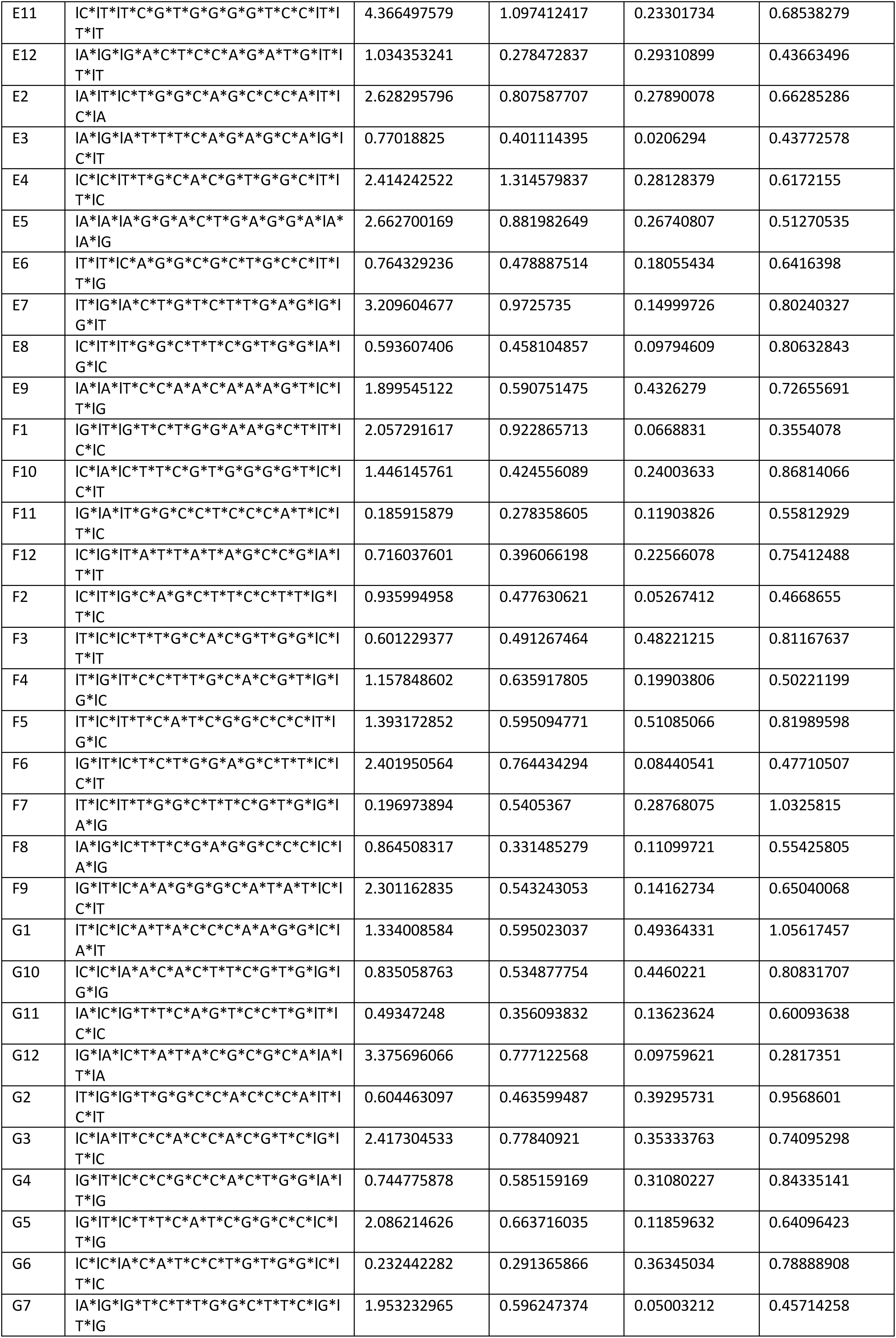
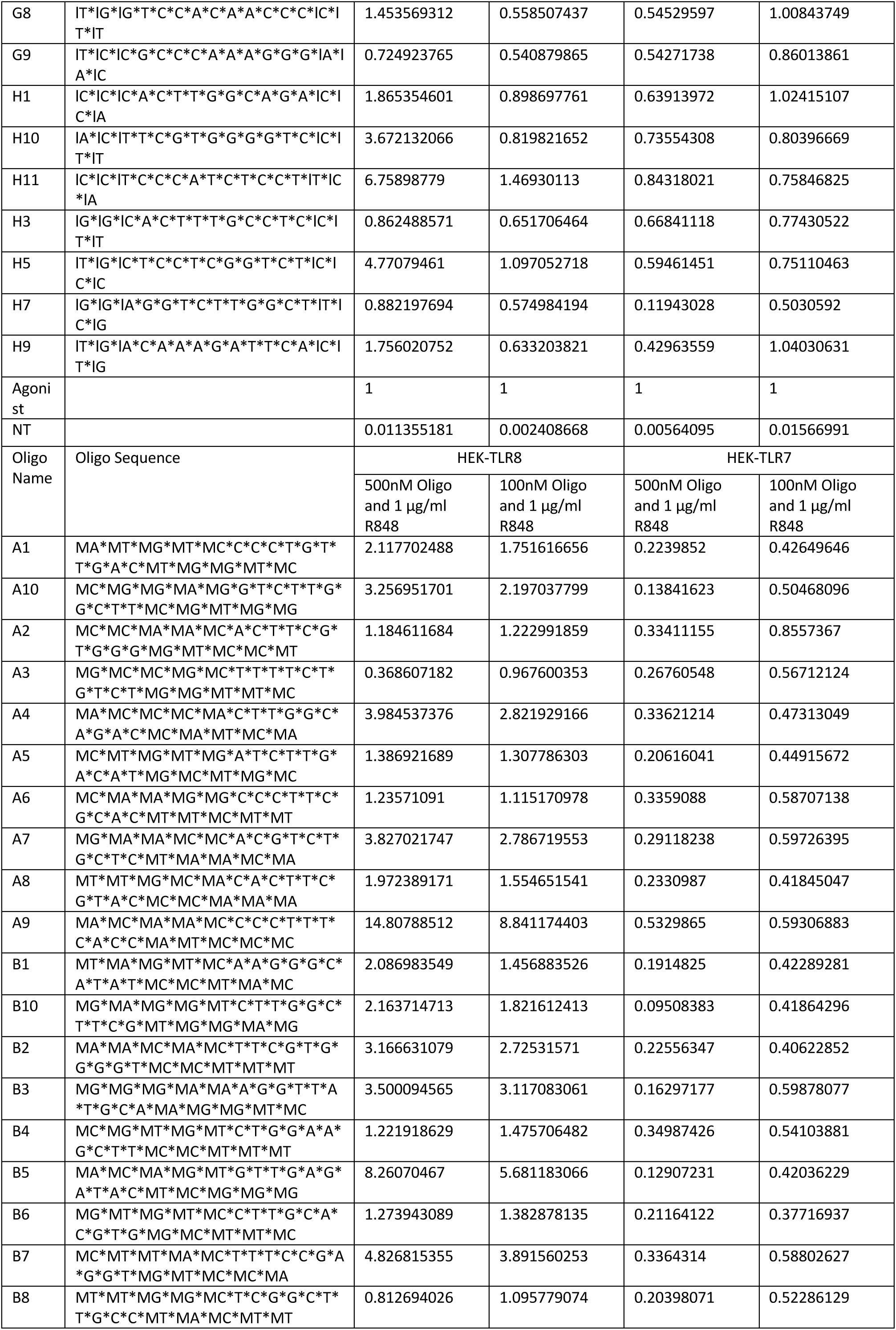
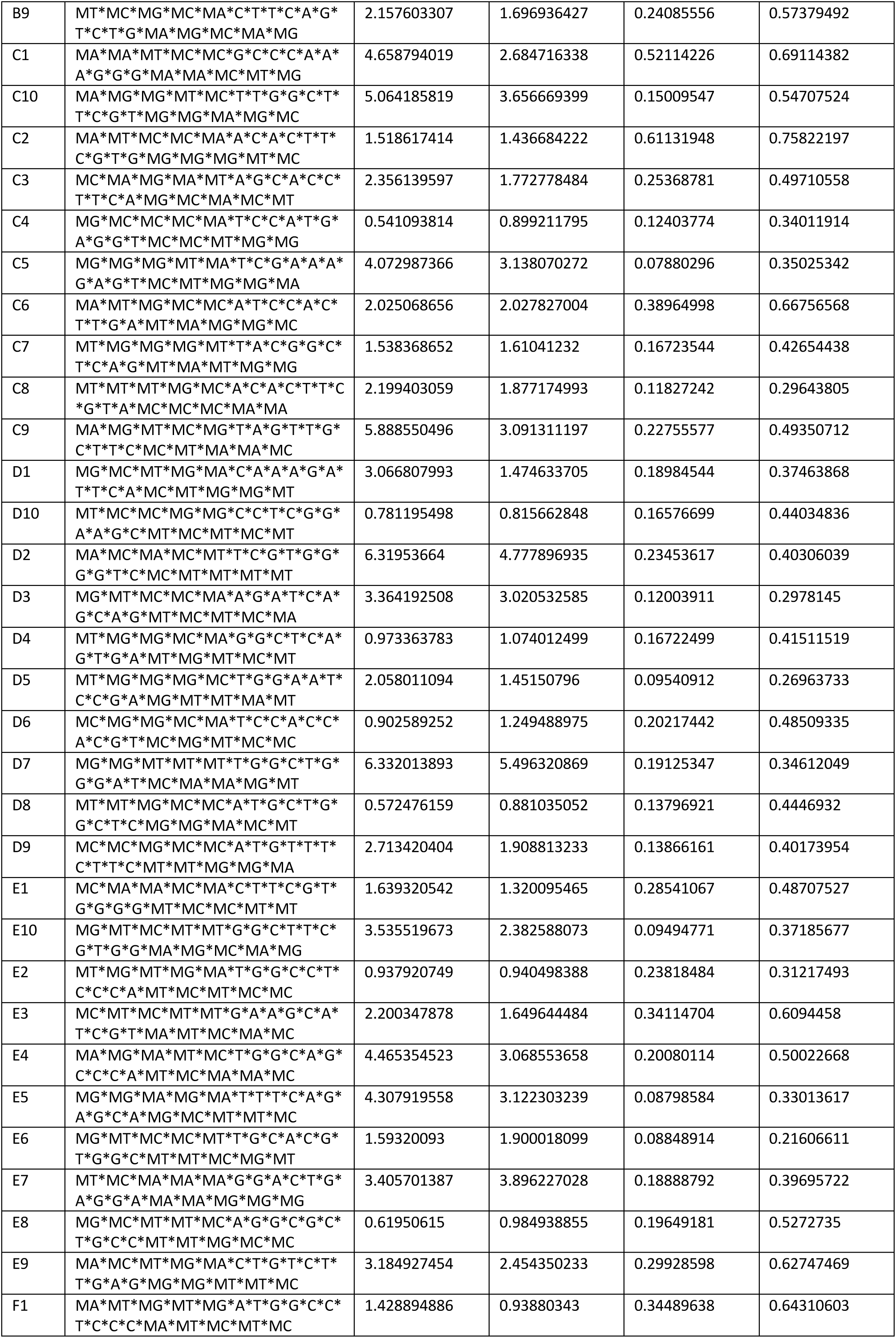
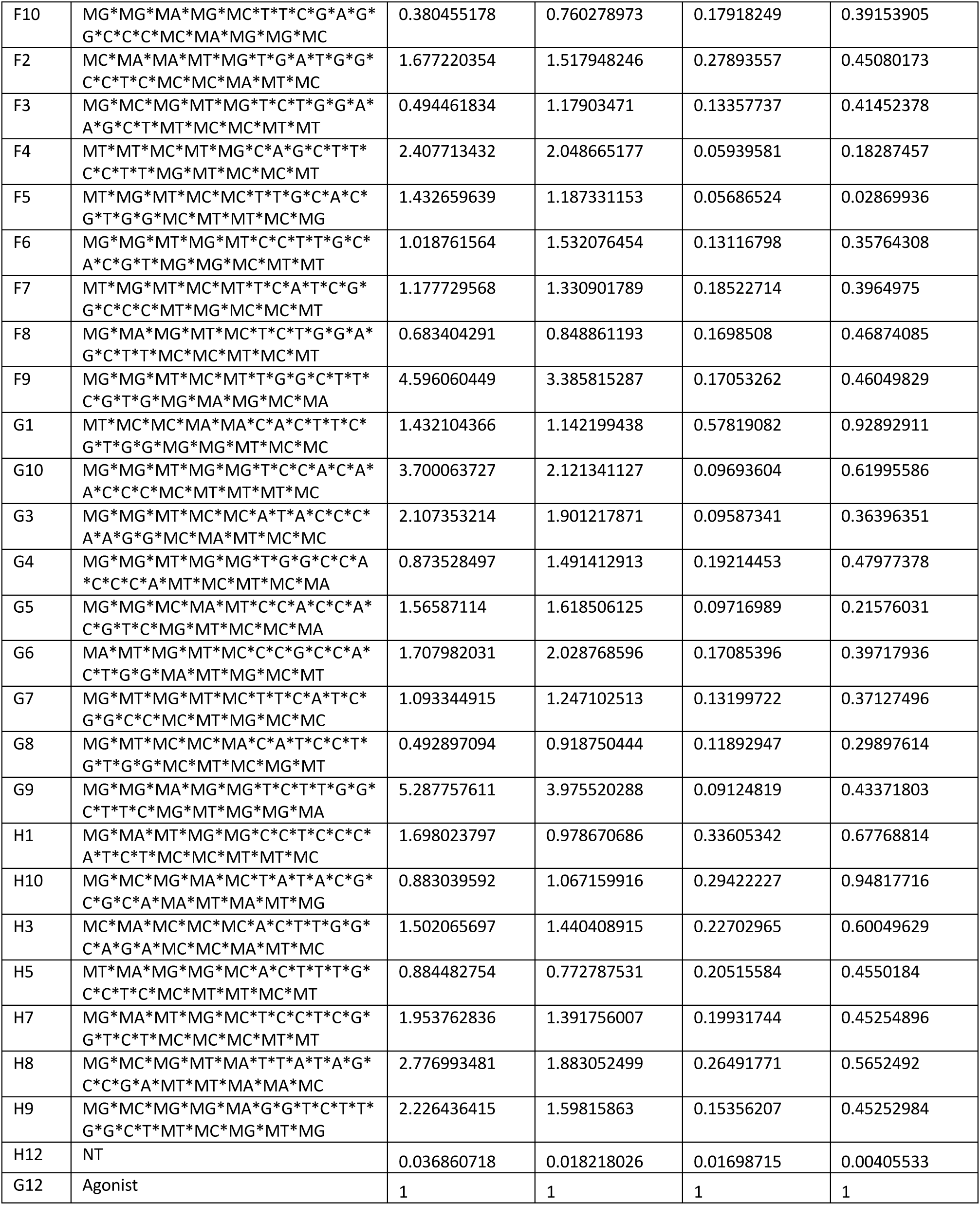
Oligonucleotides screen data. Oligonucleotides (all in 5’-3’) used in this study are labelled as: 2′-OMe is mX, DNA is either dX (for 2 and 3-mer) or UPPERCASE alone (for 16 and 20-mer), RNA is rX, LNA is I, 2’-MOE is M, and phosphorothioate inter-nucleotide linkages are denoted with a *. All values were background corrected and reported to agonist only condition are provided. The concentrations provided indicate the dose of the oligo and the agonist used in the screens.

**Table S2.**
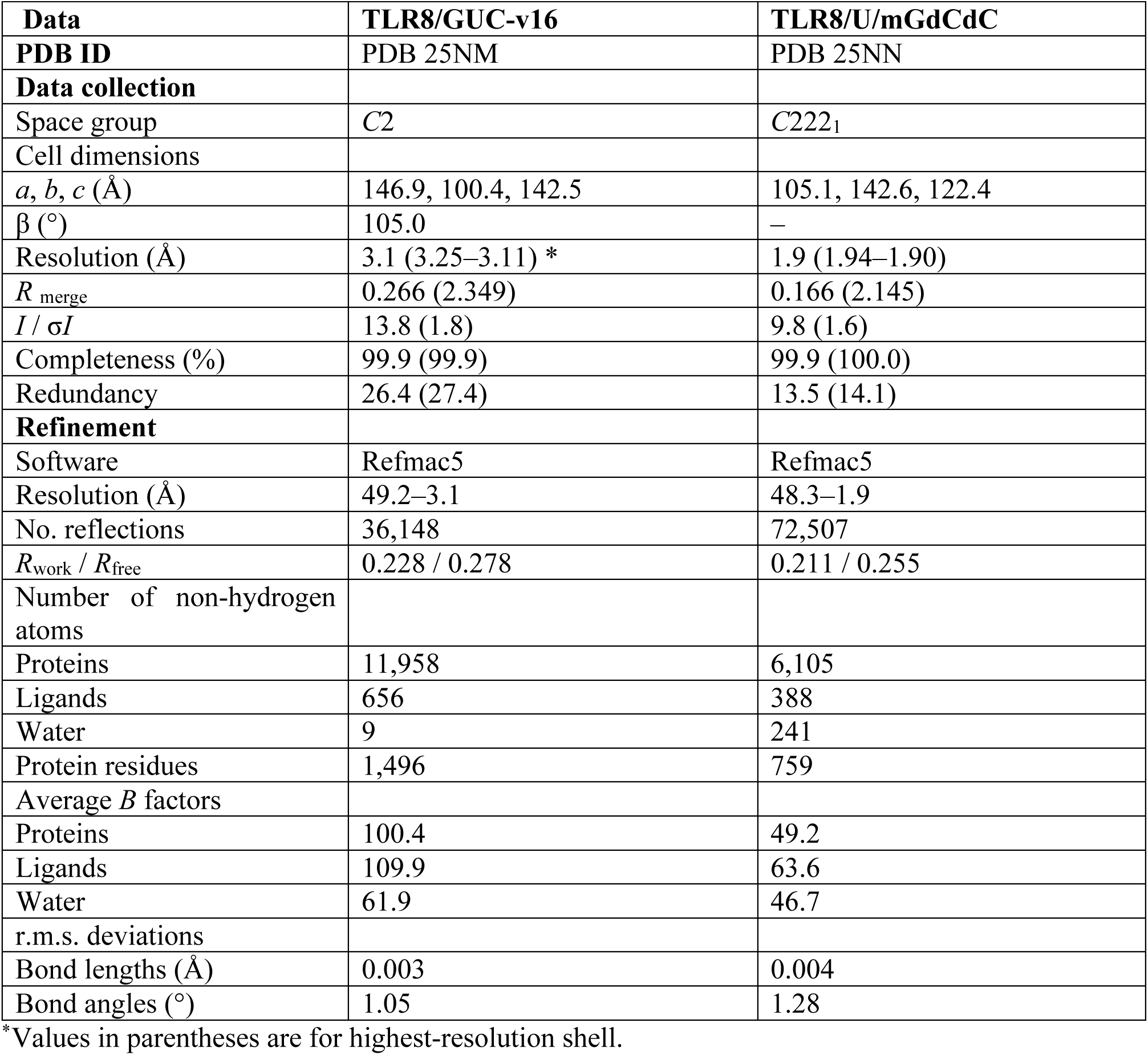
X-ray data collection and refinement statistics.

| **Data** | **TLR8/GUC-v16** | **TLR8/U/mGdCdC** |
| --- | --- | --- |
| **PDB ID** | PDB 25NM | PDB 25NN |
| **Data collection** |  |  |
| Space group | *C*2 | *C*222<sub>1</sub> |
| Cell dimensions |  |  |
| *a*, *b*, *c* (Å) | 146.9, 100.4, 142.5 | 105.1, 142.6, 122.4 |
| $\beta$ (°) | 105.0 | – |
| Resolution (Å) | 3.1 (3.25–3.11) * | 1.9 (1.94–1.90) |
| *R*<sub>merge</sub> | 0.266 (2.349) | 0.166 (2.145) |
| *I* / $\sigma I$ | 13.8 (1.8) | 9.8 (1.6) |
| Completeness (%) | 99.9 (99.9) | 99.9 (100.0) |
| Redundancy | 26.4 (27.4) | 13.5 (14.1) |
| **Refinement** |  |  |
| Software | Refmac5 | Refmac5 |
| Resolution (Å) | 49.2–3.1 | 48.3–1.9 |
| No. reflections | 36,148 | 72,507 |
| *R*<sub>work</sub> / *R*<sub>free</sub> | 0.228 / 0.278 | 0.211 / 0.255 |
| Number of non-hydrogen atoms |  |  |
| Proteins | 11,958 | 6,105 |
| Ligands | 656 | 388 |
| Water | 9 | 241 |
| Protein residues | 1,496 | 759 |
| Average *B* factors |  |  |
| Proteins | 100.4 | 49.2 |
| Ligands | 109.9 | 63.6 |
| Water | 61.9 | 46.7 |
| r.m.s. deviations |  |  |
| Bond lengths (Å) | 0.003 | 0.004 |
| Bond angles (°) | 1.05 | 1.28 |

\*Values in parentheses are for highest-resolution shell.

**Supplementary Table S3:** SPR analyses.

(a) Affinity ( $K_D$ ) values for SPR runs with wild type TLR8 and oligonucleotides (Fig. 3).
| **Oligo** | **Affinity Values ($\mu\text{M}$)** |  |  |  |
| --- | --- | --- | --- | --- |
| mG<sub>m</sub>AdG | $3.2 \pm 0.6$ | $2.4 \pm 0.2$ | $2.5 \pm 0.3$ | $2.4 \pm 0.3$ |
| mI<sub>m</sub>AdG | $0.9 \pm 0.2$ | $0.7 \pm 0.1$ | $1 \pm 0.2$ | $0.9 \pm 0.2$ |
| GCC-v4 | $11 \pm 0.7$ | $12 \pm 0.9$ | $11 \pm 0.6$ | $11 \pm 0.7$ |
| GUC-v13 | $300 \pm 30$ | $290 \pm 20$ | $320 \pm 20$ | - |
| GUC-v16 | $14 \pm 2$ | $13 \pm 2$ | $11 \pm 4$ | $14 \pm 4$ |

(b) Affinity ( $K_D$ ) values for SPR runs with wild type TLR8 or TLR8 F495S mutant and GUC-v16 (Fig. 3).
| **GUC-v16** | **Affinity Values ($\mu\text{M}$)** |  |  |
| --- | --- | --- | --- |
| TLR8 WT | $14 \pm 1$ | $13 \pm 1$ | $13 \pm 0.6$ |
| TLR8 F495S | No binding observed |  |  |
| **GCC-V4** | **Affinity Values ($\mu\text{M}$)** |  |  |
| TLR8 WT | $11 \pm 0.6$ | $12 \pm 0.5$ | $13 \pm 0.6$ |
| TLR8 F495S | $8 \pm 1$ | $9 \pm 2$ | $1 \pm 3$ |

(c) Affinity ( $K_D$ ) values for SPR runs with wild type mmTLR7 or mmTLR8 F507S mutant and GUC-v16 (Fig. 3).
| **GUC-V16** | **Affinity Values ($\mu\text{M}$)** |  |  |
| --- | --- | --- | --- |
| TLR7 WT | $1.9 \pm 0.9$ | $2 \pm 0.9$ | $1.7 \pm 0.8$ |
| TLR7-F507S | Weak binding observed |  |  |
| **Resiquimod** | **Affinity Values ($\mu\text{M}$)** |  |  |
| TLR7 WT | $4 \pm 0.8$ | $4.3 \pm 0.8$ | $4.4 \pm 1$ |
| TLR7 F507S | $3.9 \pm 0.8$ | $5 \pm 1$ | $5.3 \pm 1$ |

**Table S4.** Cryo-EM data collection, refinement and validation statistics.

| Data | TLR7/GUC-v16 |
| --- | --- |
| **EMDB ID** | EMD-80227 |
| **PDB ID** | 25NO |
| **Data collection and processing** |  |
| Microscope | Titan Krios G4 |
| Voltage (kV) | 300 |
| Detector | K3 |
| Magnification | 105,000 |
| Pixel size (Å) | 0.83 |
| Total exposure (e<sup>-</sup>/Å<sup>2</sup>) / Frames per movie | ~48 / 48 |
| Total movie stacks | 3,602 |
| Software | RELION v4.0.1 |
| Final particle images (no.) | 169,275 |
| Symmetry imposed | C2 |
| Map resolution (Å) | 3.0 |
| FSC threshold | 0.143 |
| **Refinement** |  |
| Software | Chimera, COOT, JLigand, Phenix |
| Model resolution (Å)<br>(Masked, FSC threshold = 0.5) | 3.2 |
| Model composition |  |
| Protein chains | 2 |
| Residues | 1510 |
| Ligands | GUC-v16 (*RR* isomer): 2<br>NAG: 2<br>BMA: 2 |
| Average B-factors (Å<sup>2</sup>) |  |
| Proteins | 133.9 |
| Ligands | 137.6 |
| R.m.s deviations |  |
| Bond lengths (Å) | 0.001 |
| Bond angles (°) | 0.373 |
| Validation |  |
| Molprobity score | 1.63 |
| Clashscore | 4.98 |
| Poor rotamers (%) | 1.7 |
| Ramachandran plot |  |
| Favored (%) | 96.8 |
| Allowed (%) | 3.2 |
| Outliers (%) | 0.0 |

